# An avidity-driven mechanism of extracellular BMP regulation by Twisted gastrulation

**DOI:** 10.1101/2024.07.08.602551

**Authors:** Gareth Moore, Raluca Revici, Lauren Forbes Beadle, Catherine Sutcliffe, Holly Birchenough, Clair Baldock, Hilary L Ashe

## Abstract

During dorsoventral patterning of bilaterian embryos, the conserved regulator Twisted gastrulation (Tsg) modulates BMP signalling by binding Chordin/Short gastrulation (Sog). Here we elucidate the mechanism by which Tsg interacts with Sog/Chordin to promote formation of the inhibitory Tsg-Sog/Chordin-BMP complex and regulate BMP signalling extracellularly. We identify and validate *in vitro* a hydrophobic interface in the Tsg C-terminal domain that binds Chordin. Mutation of this epitope in *Drosophila* Tsg (Tsg^L100A^) results in an unexpectedly mild perturbation to embryonic BMP gradient formation. We show that a protosome-specific Tsg C-terminal extension also binds Sog, and the presence of this second binding site allows partial rescue of Sog interaction with Tsg^L100A^ in the presence of BMP. Consistent with this, a truncated Tsg protein lacking both Sog binding regions is unable to support BMP gradient formation *in vivo*. As our data show that disruption of either Sog binding site in Tsg, but not both, can be overcome by Tsg-BMP and Sog-BMP interactions, we present a new avidity-driven mechanism of BMP gradient formation that will be relevant to a broad range of developmental contexts.

**Summary statement:** Identification and mutational analysis of the binding epitopes that mediate Twisted gastrulation interaction with Short gastrulation/Chordin reveals an avidity-based model of embryonic BMP gradient formation.

## Introduction

The Bone Morphogenetic Proteins (BMPs) are a major family of signalling molecules, belonging to the TGF-β superfamily. BMPs are involved in the development and homeostasis of many organs including the skeletal (Salazar et al., 2016) and cardiovascular (Cai et al., 2012) systems. Preceding this, BMP signalling plays a conserved role in embryonic dorsal-ventral axis patterning in Bilaterians (Mörsdorf et al., 2024). In the *Drosophila* embryo, a heterodimer of the BMP ligands Decapentaplegic (Dpp) and Screw (Scw) patterns the dorsal ectoderm (Akiyama et al., 2024). *dpp* is uniformly expressed in the dorsal ectoderm and *scw* is ubiquitously expressed (Akiyama et al., 2024). However, the Dpp/Scw heterodimer is rapidly redistributed forming a BMP activity gradient in dorsal ectoderm, peaking at the dorsal midline where the amnioserosa tissue is specified (Ferguson and Anderson, 1992a; Shimmi et al., 2005; Wang and Ferguson, 2005). Distinct thresholds of Dpp/Scw activity have been identified (Ashe et al., 2000) and the gradient is decoded by controlling the rate of promoter activation of BMP target genes (Hoppe et al., 2020).

This BMP activity gradient is established through the concerted action of a set of extracellular regulators. The antagonist *sog* is expressed in ventrolateral regions and diffuses dorsally where it binds Dpp/Scw, facilitated by collagen IV which acts as a scaffold for complex assembly (Wang et al., 2008). The Sog-Dpp/Scw complex is released from collagen IV upon the binding of Tsg and this ternary Tsg-Sog-Dpp/Scw complex diffuses dorsally (Sawala et al., 2012; Wang et al., 2008). In this ternary complex, Dpp/Scw is prevented from binding its cognate BMP receptors by Tsg and Sog. Both Tsg and the Type I receptor directly bind to BMP’s ‘wrist’ epitope (Kirsch et al., 2000a; Malinauskas et al., 2024), whereas Sog and the Type II BMP receptor bind to the BMP ‘knuckle’ epitope (Marqués et al., 1997; Piccolo et al., 1996; Zhang et al., 2007). The Dpp/Scw ligand is released from the ternary complex by the metalloprotease Tolloid (Tld), which cleaves Sog (Marqués et al., 1997). In dorsolateral regions, where Sog levels are higher, the liberated ligand is rebound by Sog and Tsg whereas at the dorsal midline, where Sog levels are lowest, the ligand binds receptors to initiate signalling (Akiyama et al., 2024; Holley et al., 1996). Through multiple rounds of binding and release, Dpp/Scw is therefore redistributed to an activity gradient peaking at the dorsal midline.

An alternative source-sink mechanism has been proposed in zebrafish, whereby BMPs diffuse dorsally towards a Chordin (vertebrate homologue of Sog) sink which is spatially restricted by Tld (Tuazon et al., 2020; Zinski et al., 2017). Despite differing mechanisms of BMP gradient formation, the presence and action of Chordin/Sog as a BMP antagonist and Tld as a BMP agonist are conserved between vertebrates and invertebrates(Mörsdorf et al., 2024). Through shuttling in the *Drosophila* embryo, Sog also acts as BMP agonist to achieve peak signalling levels at long range (Ashe and Levine, 1999). Tsg also has both agonistic and antagonistic functions. Tsg acts as an antagonist by increasing Chordin/Sog binding of BMP through the formation of the ternary inhibitory complex (Chang et al., 2001; Larraín et al., 2001; Ross et al., 2001; Shimmi et al., 2005). However, Tsg also increases the rate of Chordin/Sog cleavage by Tld to subsequently release the ligand from the inhibitory complex and therefore acts as an agonist (Larraín et al., 2001; Scott et al., 2001; Shimmi and O’Connor, 2003; Xie and Fisher, 2005).

The recent crystal structure of Tsg reveals two discretely folded domains joined by a short, flexible linker (Malinauskas et al., 2024). Tsg binds BMP ligands via its N-terminal domain (NTD) and abrogation of this interaction *in vivo* ablates BMP gradient formation in the *Drosophila* embryo (Malinauskas et al., 2024; Oelgeschläger et al., 2000). Tsg binds Chordin/Sog via its C-terminal domain (CTD) (Oelgeschläger et al., 2003). This interaction is mediated through the von Willebrand Factor type C (vWC) domains of the Chordin family of antagonists; the Chordin vWC1, vWC3, and vWC4 and Chordin-like 2 (CHRDL2) vWC1 and vWC3 domains (Fujisawa et al., 2009; Troilo et al., 2016; Zhang et al., 2007). However, the molecular mechanism by which Tsg binds Chordin/Sog and promotes ternary complex formation is so far undetermined.

Here, we combine AlphaFold predictions, *in vitro* binding studies, cell-based assays and analyses of mutant phenotypes in *Drosophila* embryos to elucidate the mechanism by which Tsg interacts with Sog/Chordin to control BMP activity. We identify two regions in *Drosophila* Tsg that bind Sog – a hydrophobic patch in the Tsg CTD that is conserved in vertebrates, and the Tsg C-terminal extension that is protostome-specific. Mutational analyses of these binding sites in cell-based assays and *in vivo* support an avidity-driven mechanism of extracellular BMP regulation.

## Results

### AlphaFold predicts a conserved Tsg-Chordin family binding mode

To identify the potential Chordin family binding surface in the Tsg CTD, we used AlphaFold-Multimer (Evans et al., 2022) to model the human Tsg-Chordin complex. AlphaFold confidently predicts the four vWC domains in addition to the four CHRD domains of Chordin (Troilo et al., 2014) and the two domains of Tsg (Malinauskas et al., 2024), with the CTD of Tsg positioned to interact with the vWC4 domain of Chordin (Fig. 1A). The interface predicted Template Modelling (ipTM) score gives a global model confidence score, weighted towards the interface residues (Evans et al., 2022; Jumper et al., 2021). Overall, the highest confidence model scored an ipTM of only 0.44, although inspection of the predicted aligned error (PAE) plot suggests a higher confidence between the relative positions of the Tsg CTD and Chordin vWC4 (Fig. 1A). Given the low confidence of the Chordin model due to long linker regions, Tsg was modelled with Chordin vWC4 alone giving a more confident prediction (ipTM of 0.65) (Fig. 1B). This proposed interface is in agreement with binding studies showing Tsg can bind to Chordin vWC4 (Zhang et al., 2007).

**Figure 1.**
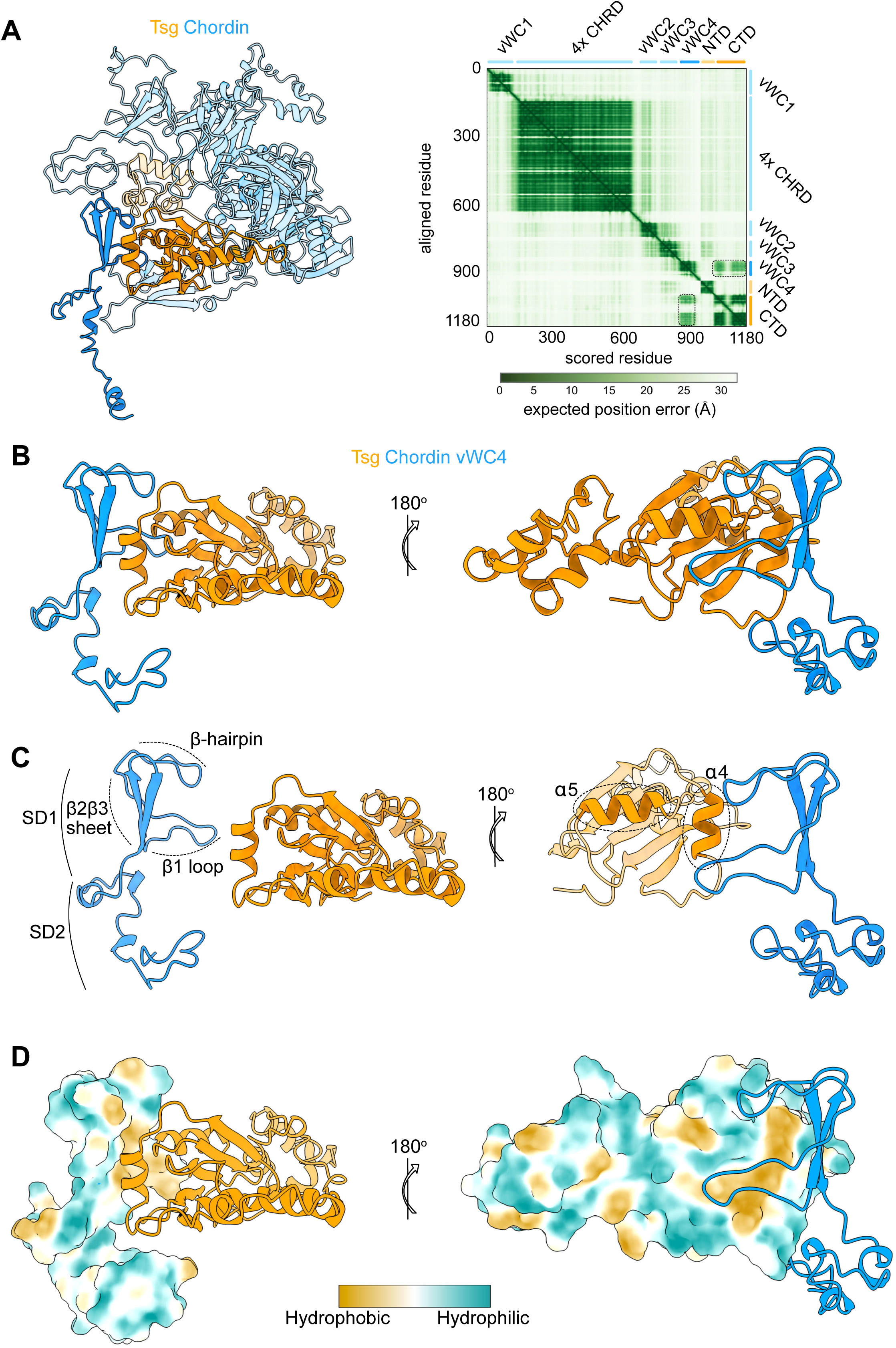
AlphaFold identifies a putative Chordin binding epitope in the Tsg CTD. (A) AlphaFold model (left) and corresponding PAE plot (right) of the human Tsg (orange) - Chordin (blue) complex with a putative interface between Tsg CTD (dark orange) and Chordin vWC4 (dark blue) as indicated by the dashed regions on the PAE plot. (B) AlphaFold model of the Tsg-Chordin vWC4 complex. (C) Model views shown in (B) but with chains separated for visibility of labelled secondary structure elements. SD1, subdomain 1; SD2, subdomain 2. (D) Tsg-Chordin vWC4 complex as in (B) with surface hydrophobicity plotted.

Inspection of the Tsg-Chordin vWC4 model shows the Tsg CTD interface is composed of two α-helices (α4 and α5) and the central β-sheet (Malinauskas et al., 2024) (Fig. 1B). This surface faces the Chordin vWC4 β-hairpin and subdomain 1 (SD1), composed of an extended loop and a two-stranded antiparallel β-sheet (Fig.1B). The loop forms an extended surface that cups the face of the Tsg CTD formed by α5 and the β-sheet, while the β2β3-strand of vWC4 sits adjacent and almost parallel to α4 of the Tsg CTD, with the β-hairpin reaching down to the intersection of Tsg CTD α4 and 5 (Fig. 1B and C). This creates a buried surface area of 823 Å^2^ between the two proteins. To gain insight into the nature of the Tsg-Chordin interaction, the surface hydrophobicity was plotted (Fig. 1D). This shows Chordin vWC4 has a central hydrophobic cavity formed by the β-hairpin (Phe874, Ala875), β1 loop (Pro887, Val889, Pro890, Pro891, Phe892, Met895) and β2β3-sheet (Ile898, Cys900, Val906, Pro907, Cys909). This faces a reciprocal hydrophobic patch on the Tsg CTD formed by α4 (Ile100, Leu103, Leu107), β-sheet (Leu114, Trp116, Phe172, Met176) and α5 (Met188).

To investigate whether Tsg could employ a common mode of interaction with the Chordin family of antagonists, human CHRDL2 and *Drosophila* Sog - Tsg complexes were modelled in AlphaFold Multimer. CHRDL2 has three vWC domains and AlphaFold models the Tsg CTD bound to vWC3 of CHRDL2 with an ipTM score of 0.48 (Fig. S1A). Modelling Tsg-CHRDL2 vWC3 specifically yields an ipTM of 0.83 and this model shows the same interface as for the Tsg-Chordin complex with the β-sheet of CHRDL2 vWC3 adjacent to Tsg CTD α4 and the loop of vWC3 cupping the Tsg CTD (Fig. S1B). AlphaFold also predicts the same mode of binding for the *Drosophila* Tsg-Sog complex (ipTM 0.5), with *Drosophila* Tsg CTD bound to vWC4 of Sog (ipTM 0.81) via the same interface (Fig. S1.C and D). Furthermore, modelling Tsg in complex with all the individual vWC domains of Chordin, CHRDL2 and Sog supports a common mode of interaction between Tsg and the Chordin family vWC domains. This suggests a conserved interaction between Tsg and the Chordin family, from flies to human.

### Conserved leucine residues mediate the Tsg-Chordin interaction

As modelling Tsg in complex with various Chordin family members suggested a common mode of interaction, we next addressed the extent to which the Tsg CTD hydrophobic patch is conserved. Alignment of the Tsg CTD from a range of species showed conservation of the hydrophobic surface centred around two highly conserved leucine residues in α4, as identified by sequence and structural alignment (Fig. 2A and Fig. S2A). As these residues, Leu103 and Leu107 in human Tsg, are highly conserved and placed at the centre of the putative Tsg- Chordin interface they were selected as candidates for mutagenesis (Fig. 2B). Each leucine residue was mutated to alanine and the effect on binding to different human Chordin family members was assessed by surface plasmon resonance (SPR). These include ΔN-Chordin, a truncated form of Chordin which is missing the N-terminal vWC1 domain but retains the C-terminal vWC2-4 domains to which Tsg binds more strongly (Troilo et al., 2016; Zhang et al., 2007), and CHRDL2 consisting of 3 vWC domains (Fig. 2C).

**Figure 2.**
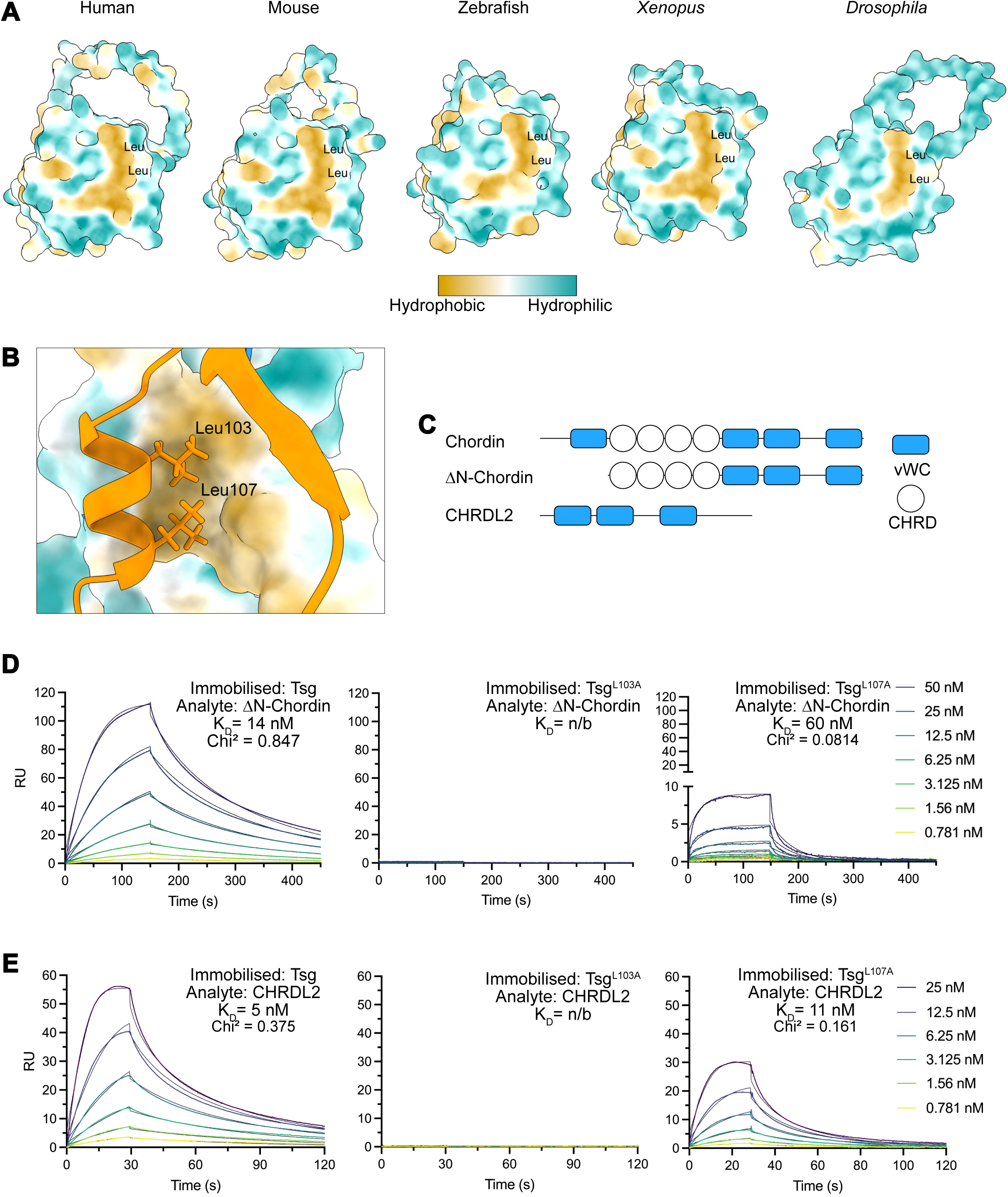
Tsg binds the Chordin family through conserved leucine residues. (A) AlphaFold models of the Tsg CTD from different species with surface hydrophobicity plotted shows a conserved hydrophobic patch centred around two conserved leucine residues. (B) Close view of the human Tsg CTD α4 helix showing positioning relative to Chordin vWC4. Conserved Leu103 and Leu107 are highlighted. (C) Domain organisation of Chordin family members and ΔN-Chordin construct. (D, E) SPR sensorgrams of ΔN-Chordin (D) and CHRDL2 (E) binding to immobilised Tsg, Tsg^L103A^, or Tsg^L107A^. Analyte was flowed over at a range of concentrations and binding kinetics calculated by fitting curves to a Langmuir 1:1 binding model. All experiments were performed in triplicate, average curves plotted with model fit in black, no binding (n/b).

Human ΔN-Chordin bound to immobilised human Tsg with high affinity (K_D_ = 14 nM), consistent with previous data (Troilo et al., 2016), but showed no interaction with Tsg^L103A^ and a weaker interaction with Tsg^L107A^ (K_D_ = 60 nM) (Fig. 2D). Similarly, CHRDL2 bound the wildtype Tsg protein tightly (K_D_= 5 nm) but showed no binding to Tsg^L103A^ and a weakened interaction with Tsg^L107A^ (K_D_ = 11 nM) (Fig. 2E). To confirm this result, we reversed the orientation of the interaction, immobilising CHRDL2 and flowing Tsg, Tsg^L103A^, or Tsg^L107A^ as the analytes. Again, in this orientation no binding of Tsg^L103A^ to CHRDL2 was observed and the affinity of Tsg^L107A^ to CHRDL2 was weaker relative to wildtype Tsg (Fig. S2B). We observed lower affinity interactions in this reversed orientation, consistent with previous binding studies of Tsg and Chordin family members (Fujisawa et al., 2009; Troilo et al., 2016; Zhang et al., 2007). To ensure the disrupted binding was not due to a destabilised Tsg CTD we assessed the thermal stability of wildtype and mutant Tsg by static light scattering (SLS). All three proteins showed remarkable thermal stability with no melting and aggregation detected from 20-95°C (Fig. S2C). This stability could be due to the high number of disulphide bonds, with 12 intramolecular disulphide bonds in the 225 residue protein (Malinauskas et al., 2024). Furthermore, neither mutation disrupted the protein secondary structure composition as assessed by circular dichroism (Fig. S2D and Table S1). Overall, these data support the proposed binding interface identified by the structural model of Tsg-Chordin and show that the conserved leucine residues are important to binding, with Leu103 particularly important in mediating the interaction. Given the consistency of this pattern in the interaction between Tsg and different Chordin family members, it suggests a common mode of binding at the molecular level.

### Embryonic Dpp/Scw gradient formation in the presence of a perturbed Tsg-Sog interaction

Given the high degree of sequence conservation of Tsg (Vilmos et al., 2001), including conservation of the key leucine residues studied above (Fig. S2A), we next tested the impact of the equivalent Leu103Ala mutation (Leu100Ala in *Drosophila* Tsg) *in vivo*. We introduced *tsg^L100A^:ALFA* sequences at the *Drosophila tsg* locus by ΦC31-mediated transgenesis in a *tsg^attP^* line (Malinauskas et al., 2024) (Fig. 3A). As the *tsg* gene is located on the X chromosome, *tsg* null males are not viable (Mason et al., 1994). Unexpectedly, given that a Sog-Tsg interaction is central to models of BMP gradient formation (Eldar et al., 2002; Ross et al., 2001; Sawala et al., 2012; Shimmi et al., 2005; Wang and Ferguson, 2005; Winstanley et al., 2015), *tsg^L100A^* males and *tsg^L100A^* homozygous females were viable. However, only a very small proportion (∼3%) of the *tsg^L100A^:ALFA* embryos hatch (Fig. S3A), indicating that *tsg^L100A^* flies are semi-viable. In addition, ∼40% of the wildtype Tsg-ALFA embryos hatch (Fig. S3A), suggesting that either the presence of the ALFA tag and/or the *mini-white* marker downstream of *tsg* following reintegration (Fig. 3A) has an effect on Tsg function or regulation.

**Figure 3.**
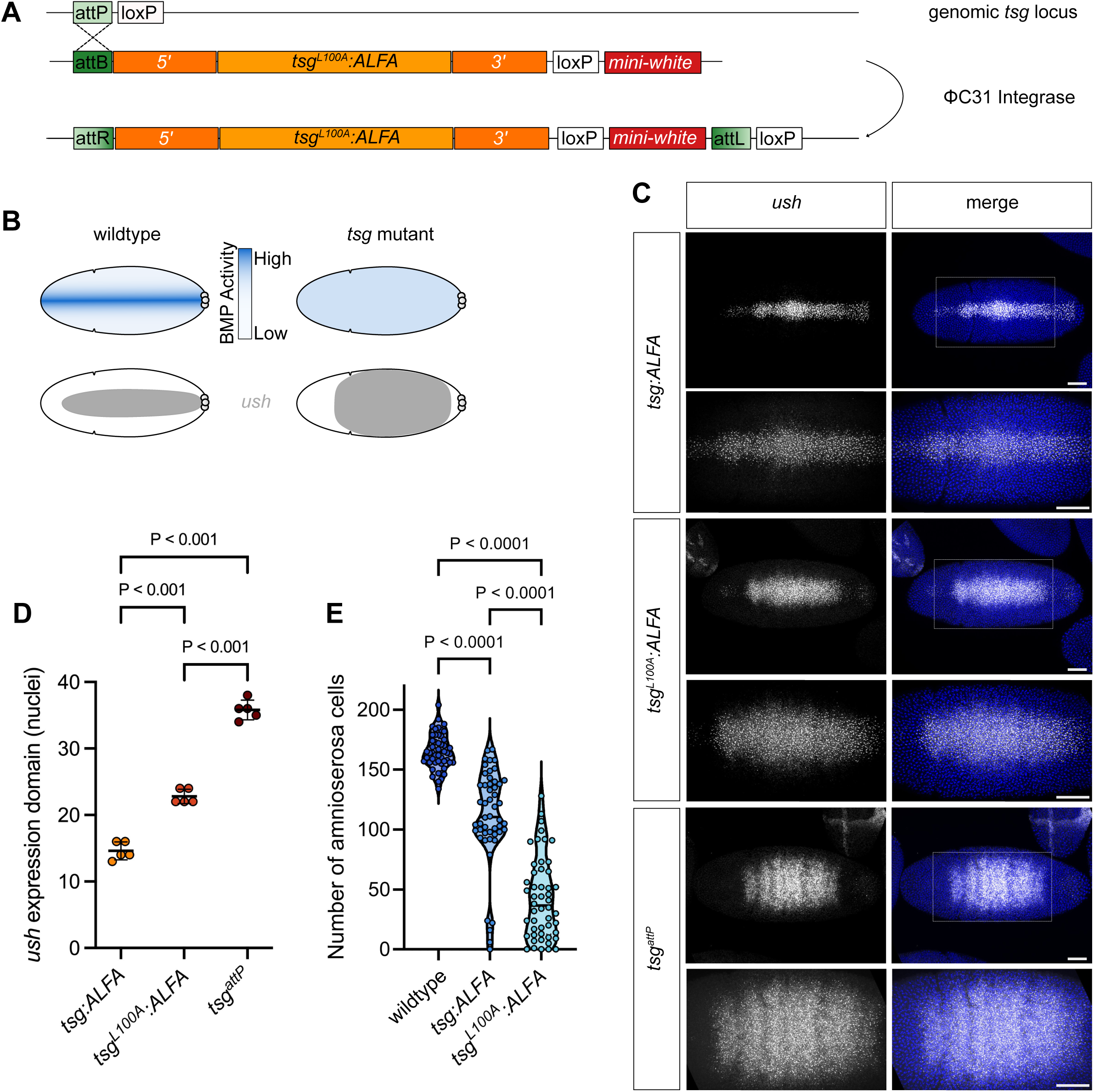
Disrupting the Tsg-Sog interaction perturbs Dpp/Scw gradient formation. (A) Schematic of the strategy used to introduce *tsg^L100A^* sequences at the *tsg* locus in *tsg^attP^* flies. (B) Cartoon of *Drosophila* embryo showing (top) the BMP activity gradient and (bottom) *ush* expression domain in wildtype and mutant *tsg* null embryos. (C) smFISH of the *ush* expression domain at 20x (top) and 40x (bottom) in *tsg:ALFA* or *tsg* mutant embryos (dorsal views). Nuclei are stained with DAPI. Scale bar = 50 μm. (D) Quantitation of the *ush* expression domain width in nuclei. One-way ANOVA with Tukey’s multiple comparisons test. n=5, mean±s.d. (E) Quantitation of amnioserosa cell number in stage 11 embryos, Kruskal-Wallis with Dunn’s multiple comparison test, n=50.

As the *tsg^L100A^* mutation is semi-viable *in vivo*, we investigated the impact on embryonic Dpp/Scw gradient formation. The BMP target gene *u-shaped* (*ush*) responds to intermediate levels of BMP activity, thereby providing a sensitive readout of gradient formation (Ashe et al., 2000) (Fig. 3B). We used single molecule fluorescence *in situ* hybridisation (smFISH) to visualise and quantitate the width of the *ush* expression (Fig. 3C). Null *tsg^attP^* embryos fail to establish a BMP activity gradient and therefore display an expanded *ush* expression domain, while reintegration of *tsg:ALFA* sequences rescues gradient formation to establish the wildtype *ush* pattern (Malinauskas et al., 2024) (Fig. 3C, D). Mutant *tsg^L100A^:ALFA* embryos display a broader *ush* expression domain than *tsg:ALFA* embryos, however this pattern is not as broad as in null *tsg^attP^* embryos where gradient formation is absent (Fig. 3C, D).

We also combined immunostaining of pMad, the activated BMP transducer, with FISH imaging of *Race*, a peak BMP target gene (Ashe et al., 2000), and *ush* in control, *tsg:ALFA* and *tsg^L100A^:ALFA* embryos. In control and *tsg:ALFA* embryos, pMad accumulation is restricted to a narrow stripe at the dorsal midline (Fig. S3B) (Dorfman and Shilo, 2001). Although there is no significant difference in the width of the pMad stripe at the dorsal midline, *Race* expression is narrower in the *tsg:ALFA* embryos compared to control embryos (Fig. S3B, C). As activation of *Race* expression is an exquisitely sensitive readout of peak BMP signalling levels (Ashe and Levine, 1999), the weaker *Race* activation in the *tsg:ALFA* embryos suggests that these embryos have a minor reduction in peak BMP signalling that is not evident from the pMad staining. There is no significant difference in the width of the *ush* expression domain in the control and *tsg:ALFA* embryos (Fig. S3B, C).

In the *tsg^L100A^:ALFA* embryos, the pMad stripe is wider than observed for *tsg:ALFA* embryos, consistent with a less refined pMad gradient (Fig. S3B, C). The mean width of the *Race* expression domain is not significantly different from the *tsg:ALFA* embryos, but there is a trend that *tsg^L100A^:ALFA* embryos have weaker *Race* expression including a higher proportion that completely fail to activate *Race* (Fig. S3C). In addition, the *ush* expression domain is significantly broader in the *tsg^L100A^:ALFA* embryos compared to the *tsg:ALFA* and control embryos (Fig. S3B, C), as observed with smFISH staining (Fig. 3C). The *ush* expression domain does not extend throughout the whole dorsal ectoderm (Fig. S3B), indicating that there are graded BMP levels in the *tsg^L100A^:ALFA* embryos. Together, these data support a shallower BMP gradient in the *tsg^L100A^:ALFA* embryos.

To assess the impact of this less refined BMP activity gradient on patterning, the number of amnioserosa cells in stage 11 embryos was determined using Hindsight staining (Fig. S3D). Quantitation shows that the number of amnioserosa cells is significantly lower and more variable in *tsg:ALFA* embryos compared to wildtype control embryos (Fig. 3E). Around half the *tsg:ALFA* embryos have less amnioserosa cells than wildtype, suggesting that the effects of the slightly weaker peak BMP signalling in the *tsg:ALFA* embryos described above (Fig. S3) are exacerbated at this later stage. The proportion of *tsg:ALFA* embryos with lower numbers of amnioserosa cells (58% have <125 amnioserosa cells) mirrors the proportion that fail to hatch (Fig. S3A). This lethality is consistent with a previous observation that embryos can only tolerate a 20% reduction in amnioserosa cell number (Gavin-Smyth et al., 2013).

*tsg^L100A^:ALFA* embryos show disrupted amnioserosa specification with significantly fewer amnioserosa cells compared to *tsg:ALFA* embryos (Fig. 3E). The spread of amnioserosa cell numbers may reflect variations in the steepness of the BMP gradient between *tsg^L100A^:ALFA* embryos, as the widths of the pMad stripe and ush expression domain also show variation (Fig. S3C). Again, the proportion of *tsg^L100A^:ALFA* embryos within the wildtype range of amnioserosa cell number (2% have >125 amnioserosa cells) is similar to the hatching rate (Fig. S3A). Together, these data suggest that limited gradient formation is established in the dorsal ectoderm of *tsg^L100A^:ALFA* embryos, but the shallower gradient is sufficient to specify a wildtype number of amnioserosa cells and permit full development in only a very small proportion of *tsg^L100A^:ALFA* embryos.

### An avidity-based model for BMP-Sog-Tsg complex formation

To understand how abrogation of the Tsg-Sog interaction could allow formation of a shallow BMP gradient, we directly tested the impact of the Leu100Ala mutation on the binding of *Drosophila* Tsg to Sog based on expression of these proteins in S2R+ cells. We performed a co-immunoprecipitation assay by mixing Tsg:ALFA or Tsg^L100A^:ALFA conditioned media with Sog conditioned media in the presence and absence of Dpp:HA/Scw:Flag conditioned media. Tsg was immobilised on anti-ALFA resin and the amount of Sog and Dpp bound was assessed by Western blot (Fig. 4A). Tsg binds Sog and this interaction is increased in the presence of Dpp/Scw (Ross et al., 2001; Shimmi et al., 2005) (Fig. 4A). Tsg^L100A^ did not bind Sog (Fig. 4A), consistent with the findings for the human proteins described above, which further supports the conserved binding mode between Tsg and the Chordin family of antagonists. However, in the presence of Dpp/Scw, we observed rescue of the Tsg-Sog interaction, albeit to a lesser extent compared to wildtype Tsg (Fig. 4A). This suggests that, upon addition of Dpp/Scw, the presence of BMP-Sog and BMP-Tsg interactions can support a Tsg-Sog interaction even though the interface harbours the L100A mutation. The ability to rescue the loss of one interaction by others is consistent with an avidity mechanism, in which ternary complex formation depends on the cooperative binding of multiple interactions (Fig. 4B). Although avidity can rescue ternary complex formation, there is less with Tsg^L100A^ compared to wildtype Tsg (Fig. 4A), which can explain the shallower BMP gradient and disrupted amnioserosa formation in *tsg^L100A^:ALFA* embryos (Fig. 3C-E).

**Figure 4.**
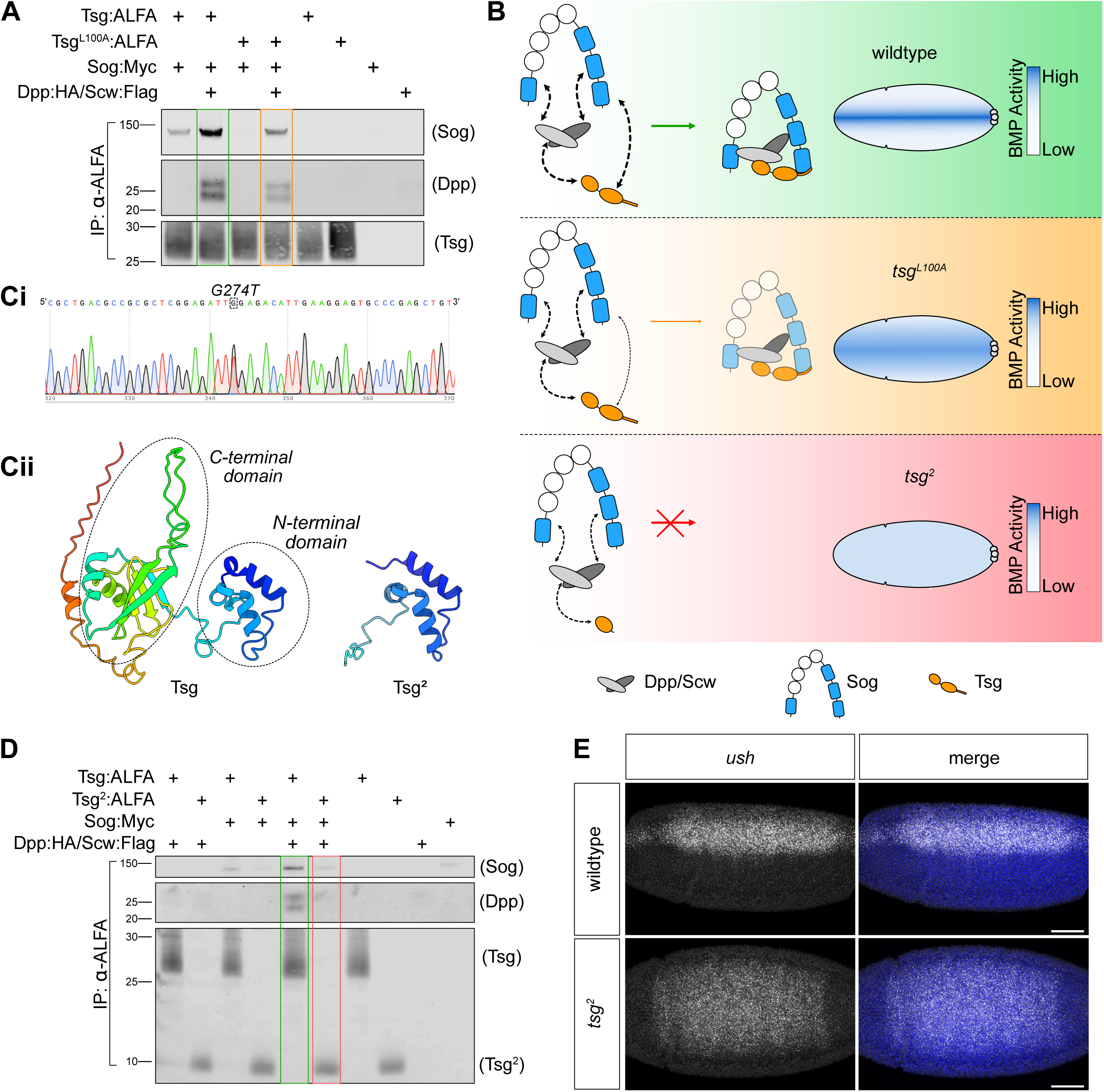
An avidity driven model of inhibitory ternary complex formation. (A) Western blot showing the amount of Sog and Dpp co-immunoprecipitated with Tsg:ALFA or Tsg^L100A^:ALFA proteins, immobilised on anti-ALFA resin. Combinations of protein conditioned media from S2R+ cells are as shown in the key. The co-immunoprecipitation assay was performed in triplicate. Green and orange boxes correspond to equivalent colours in B. (B) Cartoon of proposed avidity model of ternary complex formation and subsequent BMP gradient pattern in wildtype and mutant contexts. The thickness of the double-headed arrows indicates the strength of the interactions. (C) Characterisation of the *tsg^2^*mutant by i) genomic sequencing identifying the point mutation and ii) visualisation of the subsequent protein truncation. (D) As in (A) but for Tsg:ALFA or Tsg^2^:ALFA proteins. Green and red boxes correspond to the equivalent colours in B. (E) *ush* FISH in wildtype or *tsg^2^*embryos (dorsal view). Nuclei are stained with DAPI. Scale bar = 50 μm.

As the avidity model proposes that strong Tsg-BMP and Sog-BMP interactions can support Tsg-Sog interaction when the interface is weakened, a prediction of the model is that complete loss of the Sog-interacting interface in Tsg would not be tolerated. To test this, we have exploited *tsg^2^*, a null allele isolated ∼30 years ago, for which the molecular basis was unknown (Mason et al., 1994; Wieschaus et al., 1984). By sequencing the *tsg^2^* allele, we show that it carries a nonsense mutation after the N-terminal (BMP-binding) domain (Fig. 4Ci) resulting in a truncated protein (Fig. 4Cii). As this protein lacks the C-terminal domain including the residues important for Sog interaction, we tested its ability to form the ternary complex. Using the S2R+ cells co-immunoprecipitation assay described above, we show that the Tsg^2^:ALFA protein cannot bind Sog, even when Dpp/Scw is present, and neither is the Tsg-Dpp/Scw interaction augmented (Fig. 4D). This is consistent with an inability of Dpp/Scw to rescue ternary complex formation through avidity as the truncated Tsg^2^ protein has no C-terminal domain for Sog to bind to, despite retaining the BMP-binding NTD (Fig. 4B). Visualisation of *ush* (Fig. 4E)*, Race* and pMad (Fig. S4) in *tsg^2^* embryos shows that there is a loss of BMP gradient formation, as reported previously (Malinauskas et al., 2024; Mason et al., 1994). These data provide a molecular explanation for the absence of a BMP gradient in *tsg^2^* embryos and their lethality (Malinauskas et al., 2024; Mason et al., 1994; Wieschaus et al., 1984). Together, these results demonstrate that mutation, but not loss, of the Sog-interacting interface in Tsg can support BMP gradient formation *in vivo* due to an avidity mechanism involving the synergistic action of all three components of the shuttling complex (Ross et al., 2001; Shimmi et al., 2005).

### A C-terminal Tsg tail sequence, which is conserved in protostomes, is required for Sog interaction and BMP gradient formation

To further interrogate avidity driven ternary complex formation, we next asked whether avidity rescues Tsg^L100A^ binding to Sog through the remaining residues in the Tsg hydrophobic interface or by a secondary Sog-binding site in Tsg. In *Drosophila* species, Tsg also has a C-terminal extension of 40 residues, termed a tail, which was found to be absent from Tsg in the vertebrate species analysed (Vilmos *et. al* 2001). As the Tsg tail is of unknown function, yet proximal to the Sog-binding CTD, we extended our study to investigate its requirement for Tsg activity. First, we performed a more detailed analysis of conservation of the Tsg tail. Alignment of 60 Tsg sequences from a broad range of species revealed a dichotomy in the C-termini of different *tsg* sequences; 32/60 species had a consistent C-terminus two residues after the final conserved cysteine, whereas the remaining 28/60 species showed some degree of C-terminal extension (Fig. S5). Phylogenetic analysis revealed that the presence or absence of a tail sequence traced back some 600 million years ago at the protostome-deuterostome split. All 28 species with an extension were protostomes and all 32 species lacking Tsg tail sequences were deuterostomes. While clearly present at the C-terminus of protostome Tsg sequences, there is little-to-no conservation at the sequence level, beyond an enrichment in acidic residues (Fig. S6A), or indeed the length of the tail.

AlphaFold does not predict any significant secondary structure in the *Drosophila* Tsg tail (Fig. S6B), and this prediction was supported by other *in silico* tools that score the Tsg tail region as likely to be an intrinsically disordered region (IDR) (Fig. S6C). In the absence of any structural information, we used the co-immunoprecipitation assay in S2R+ cells to test whether deleting the tail affected Tsg’s interaction with Sog or Dpp/Scw heterodimers. We expressed Tsg^Δtail^:ALFA, a Tsg truncation (Met1-Glu208) that removes the tail sequence (Asp209-Ser249) two residues after the final cysteine as in deuterostome sequences. Wildtype or truncated Tsg:ALFA conditioned media were mixed with Dpp:HA-Scw:Flag and/or Sog:Myc conditioned media before immobilising Tsg:ALFA with anti-ALFA resin and detecting bound protein by Western blot (Fig. 5A). While the truncated Tsg^Δtail^ protein bound Dpp equally to wildtype Tsg, Tsg^Δtail^ did not bind Sog (Fig. 5A), demonstrating that the Tsg tail is also important for Sog interaction. However, in the presence of Dpp/Scw, the Tsg^Δtail^ binding to Sog was partially rescued, to a lower level of binding than the wildtype Tsg-Sog interaction in the presence of the ligand (Fig. 5A). This mirrors the finding that the Tsg^L100A^-Sog interaction can be partially rescued in the presence of Dpp/Scw (Fig. 4A).

**Figure 5.**
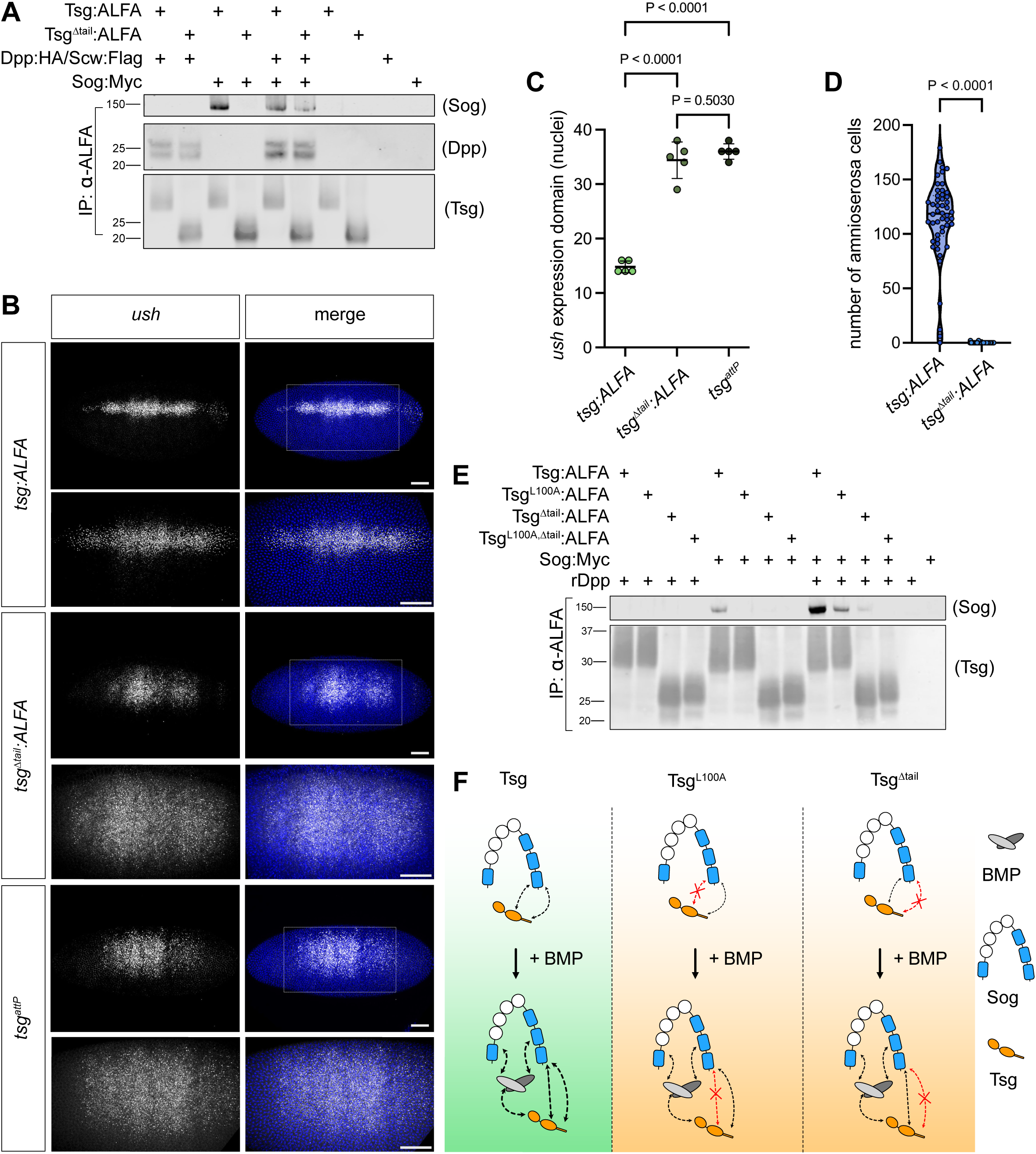
The C-terminal Tsg tail sequence is essential to Tsg function. (A) Western blot showing the co-immunoprecipitation of Dpp and Sog with Tsg:ALFA or truncated Tsg^Δtail^:ALFA. Protein combinations, from conditioned S2R+ cell media, are shown in the key. The co-immunoprecipitation assay was performed in triplicate. (B) smFISH of *ush* expression in *tsg:ALFA*, *tsg^attP^*, and *tsg^Δtail^:ALFA* embryos, dorsal views, nuclei are stained with DAPI. Scale bars = 50 μm. (C) Plot of *ush* expression domain width in nuclei, one-way ANOVA with Tukey’s multiple comparisons test, n=5, mean±s.d. (D) Number of amnioserosa cells in stage 11 embryos of the indicated genotypes. Mann Whitney test, n=50. (E) Western blot showing the co-immunoprecipitation of Sog with Tsg:ALFA, Tsg^L100A^:ALFA, Tsg^Δtail^:ALFA, or Tsg^L100A/Δtail^:ALFA. The co-immunoprecipitation assay was performed in triplicate. (F) Cartoon diagram of additive Sog-binding sites in Tsg and the mechanism of partial ternary complex rescue through avidity. Weight of dashed arrows indicates interaction strength.

As the Tsg^Δtail^ protein can still form a ternary complex (Fig. 5A), we tested whether, like Tsg^L100A^:ALFA, Tsg^Δtail^:ALFA could support BMP gradient formation *in vivo*. *tsg^Δtail^:ALFA* sequences were reintegrated into the *tsg^attP^* locus, which resulted in male lethality with 100% penetrance. To investigate the effect on BMP gradient formation, we first visualised expression of *ush* (Fig. 5B). Embryos containing the truncated *tsg* sequences show a broad expansion of the *ush* expression domain as in null *tsg^attP^* embryos (Fig. 5C), although the degree of expansion along the anterior-posterior axis appears more variable in the *tsg^Δtail^:ALFA* embryos (Fig. 5B, S7A). pMad immunostaining shows that the *tsg^Δtail^:ALFA* embryos fail to accumulate pMad at the dorsal midline, whereas FISH reveals that *Race* expression in the presumptive amnioserosa is lost and *ush* expression is expanded in the dorsal ectoderm (Fig. S7A). These phenotypes are characteristic of a lack of BMP gradient formation in the *tsg^Δtail^:ALFA* embryos. Quantitation of the number of amnioserosa cells in *tsg^Δtail^:ALFA* embryos revealed a failure to specify amnioserosa (Fig. 5D, Fig. S7B), consistent with a loss of peak BMP signalling (Mason et al., 1994; Wang and Ferguson, 2005). This demonstrates that the Tsg tail contributes an essential function during BMP gradient formation. Moreover, the lack of a BMP gradient and amnioserosa specification explains the loss of viability in *tsg^Δtail^:ALFA* males.

Tsg^L100A^:ALFA can weakly support BMP gradient formation *in vivo*, whereas Tsg^Δtail^:ALFA cannot, even though in the co-immunoprecipitation assays both Tsg^L100A^:ALFA and Tsg^Δtail^:ALFA form ternary complex in the presence of Dpp/Scw (Fig. 4A, 5A). One potential explanation is that there is weaker rescue of ternary complex formation by BMP with Tsg^Δtail^:ALFA. To address this, we directly compared ternary complex formation using S2R+ co-immunoprecipitation assays. For these experiments, instead of Dpp/Scw from conditioned media, we used recombinant Dpp which can be added at higher concentration to give a more robust enhancement of ternary complex formation. As before, both Tsg^L100A^:ALFA and Tsg^Δtail^:ALFA only bind Sog in the presence of Dpp, to an overall lower level than with wildtype Tsg (Fig. 5E). There was some variability in the amount of ternary complex formed with Tsg^Δtail^, but generally less ternary complex was formed than with Tsg^L100A^:ALFA (Fig. 5E). This suggests that the inability of *tsg^Δtail^:ALFA* embryos to form a BMP gradient is either due to insufficient ternary complex formation or the Tsg tail also has a second function (see Discussion).

We also generated a double Tsg^L100A,Δtail^:ALFA mutant to test whether the hydrophobic interface and tail function additively in relation to Sog interaction. Binding of the double Tsg^L100A, Δtail^:ALFA mutant to Sog is not rescued in the presence of Dpp (Fig. 5E), similar to the lack of rescue observed for the Tsg^2^ mutant (Fig. 4D). This reveals that the conserved hydrophobic domain and tail region in Tsg contribute additively to Sog binding, and the increase in binding of either Tsg^L100A^:ALFA or Tsg^Δtai^:ALFA to Sog in the presence of Dpp is mediated through the other, unperturbed, Sog binding site in Tsg (Fig. 5F). Together, these results show that despite a lack of sequence or structural conservation, the protostome-specific Tsg tail is necessary for the full action of the protein, with specificity to Sog but not Dpp/Scw binding, during dorsoventral patterning in *Drosophila* embryogenesis.

## Discussion

In this study we used *in silico* structural modelling with AlphaFold to identify a binding site in Tsg for Chordin family members, which is conserved from flies to humans. Modelling proposed a conserved binding mode between the Tsg CTD and Chordin family vWC domains, consistent with previous binding studies showing that Tsg binds the Chordin family via its CTD (Malinauskas et al., 2024; Oelgeschläger et al., 2003). The newly identified interface has a hydrophobic core centred around a pair of conserved leucine residues, Leu103 and Leu107 in human Tsg. The reciprocal hydrophobic surface in the Chordin vWC domain is formed by an extended loop and β-sheet that comprises SD1 of the vWC domain. Mutation to residues in this putative interface in CHRDL2 vWC domains reduced their affinity for Tsg (Fujisawa et al., 2009), consistent with the proposed binding interface identified here. The extended SD1 loop region of Chordin family vWC domains has been proposed to be a major determinant of binding BMPs and Tsg (Fujisawa et al., 2009; Xu et al., 2017). This is in line with the AlphaFold models of Tsg-Chordin family antagonists presented here. Analysis of vWC domains across Chordin family members shows those containing an extended loop region in SD1 bind both Tsg and BMP. However, vWC2 of both Chordin and CHRDL2, with a short linker, does not bind Tsg or BMP (Fujisawa et al., 2009; Troilo et al., 2016; Zhang et al., 2007). This supports the proposal that the SD1 loop mediates binding of Chordin family vWC domains for a general model of binding to Tsg. Domain specific sequence differences in this SD1 loop could then explain the differential binding ability of Chordin family vWC domains.

In addition to this highly conserved hydrophobic interface, we identified the Tsg tail as a second region important for Sog interaction. The presence of the Tsg tail in protostomes but not deuterostomes is striking. A notable evolutionary difference in extracellular BMP regulation between protostomes and deuterostomes is the ligand-dependent activity of Tld. In *Drosophila*, Sog is only cleaved by Tld when bound to a BMP ligand (Marqués et al., 1997; Peluso et al., 2011), whereas vertebrate Tld is able to process Chordin without a BMP ligand (Piccolo et al., 1997). Therefore, we speculate that the Tsg tail could relate to the BMP-dependence of Sog cleavage in protostomes. Alternatively, the evolutionary divergence in Tsg with respect to the tail could relate to the mechanism and dynamics of gradient formation. In the protostome *Drosophila*, the steep BMP activity gradient is rapidly refined through a shuttling mechanism and positive feedback (Ferguson and Anderson, 1992; Shimmi et al., 2005; Wang and Ferguson, 2005). In the deuterostome *Danio rerio*, the BMP gradient evolves over greater length and time scales to a comparatively shallower gradient through a source-sink mechanism (Tuazon et al., 2020; Zinski et al., 2017). These different dynamics could mean source-sink based systems do not require the tail to strengthen Tsg-Chordin interactions, and therefore ternary complex formation, as rapidly as in flies.

We propose that the disordered negatively charged Tsg tail could bind to the positively charged vWC domain of Sog, consistent with negatively charged IDRs being able to stay disordered and bind to positively charged folded domains (Bugge et al., 2025). Our binding data show that, as observed with Tsg^L100A^, Tsg^Δtail^ binding of Sog is partially rescued in the presence Dpp/Scw. However, while introduction of the L100A mutation *in vivo* can support formation of a shallow BMP gradient and viability, truncation of the Tsg tail *in vivo* results in a loss of the BMP gradient and lethality. The stronger phenotype observed with the *tsg^Δtail^* mutant could reflect slightly weaker ternary complex formation, which falls below a level sufficient for BMP gradient formation. Alternatively, since Tsg^Δtail^ can form some ternary complex, it is possible that the Tsg tail is needed for more than just mediating Tsg-Sog interaction. Possibilities include, firstly, the Tsg tail could promote Sog cleavage in a BMP-dependent manner, as mentioned above. This could be mediated by an interaction with Tld or by promoting a conformational change in Sog (Troilo et al., 2016). Secondly, the acidic residue-rich Tsg tail may be required for Tsg’s function in releasing ternary Tsg-Sog-Dpp/Scw shuttling complexes from a collagen IV scaffold (Wang et al., 2008), given that Sog binds collagen IV via basic residues (Sawala et al., 2012). Thirdly, the tail may be needed for another aspect of Tsg action, such as movement of Tsg which is highly diffusive (Mason et al., 1997), for example by increasing solubility (Fig. S6D,E). Investigation into the molecular action of the Tsg tail offers an interesting direction for future research.

Based on our mutational analyses, we propose that the simplest hypothesis for Tsg promoting ternary complex formation is through an avidity mechanism, where binding of one epitope makes a second interaction more likely. Firstly, we show that in the presence of weakened Sog-Tsg interactions, due to the Tsg^L100A^ or Tsg^Δtail^ mutations, ternary complex formation can be rescued by Sog-BMP and Tsg-BMP interactions. Secondly, we demonstrate that the Tsg^2^ truncation, which removes both the C-terminal domain and tail, cannot promote ternary complex formation or support BMP gradient formation *in vivo*. Due to the truncation of the Sog interacting region in Tsg, the remaining BMP-Sog and BMP-Tsg interactions become decoupled and are themselves insufficient to support ternary complex formation. Thirdly, as Tsg^2^-BMP interaction is not enhanced in the presence of Sog, this shows all three interactions are necessary to increase the binding of any of them, consistent with an avidity mechanism. Finally, using the S2R+ based assay, we demonstrate that the Tsg^L100A, Δtail^ double mutant protein is unable to form any ternary complex. This finer dissection supports that the defect in the Tsg^2^ protein is loss of the hydrophobic interface and tail that normally function additively to increase ternary complex formation through avidity, but can each also weakly support ternary complex independently.

In terms of the molecular interactions supporting avidity, Chordin family vWC domains bind BMP ligands at the type II receptor knuckle epitope (Zhang et al., 2007) so, despite containing four vWC domains, only two vWC domains of Sog/Chordin can bind one BMP dimer at once. Tsg binds the BMP ligand at the wrist (type I receptor) epitope through its NTD (Malinauskas et al., 2024; Zhang et al., 2007). By simultaneously binding the BMP wrist epitope, through its NTD, and Sog/Chordin, through its CTD, Tsg provides an additional link between Sog/Chordin and the BMP ligand. Interaction at one binding site could then aid subsequent interaction by bringing the remaining epitopes closer, thus promoting interaction via an avidity mechanism (Fig. 4B). Such a mechanism of Tsg action is supported by the observation that the linker region is of conserved length, suggesting that the spacing of the NTD and CTD is key to function (Malinauskas et al., 2024). Furthermore, abrogation of the Tsg-Dpp/Scw interaction leads to a complete loss of ternary complex formation and gradient establishment in the *Drosophila* embryo (Malinauskas et al., 2024). In contrast to this complete abrogation, the semi-viability of the *tsg^L100A^:ALFA* embryos allowed us to uncover the molecular evidence supporting an avidity-driven mechanism of ternary complex formation. As ternary complex formation in *tsg^L100A^:ALFA* embryos is rescued due to the presence of the protostome-specific Tsg tail, the avidity mechanism only became apparent through the combined study of the human and *Drosophila* proteins.

Avidity of Tsg-Sog/Chordin binding the BMP ligand would mirror BMP receptor complex formation, which is thought to assemble through avidity-driven complexing (Groppe et al., 2008). In the context of BMP gradient formation in the early *Drosophila* embryo, such avidity of Tsg-Sog/Chordin binding to BMP would be important. Quantitative models of Dpp/Scw shuttling require Sog to bind the ligand in the low pM range (Eldar et al., 2002; Mizutani et al., 2005; Umulis et al., 2006), while binding studies with the vertebrate homologues suggest the interaction is in low nM range (Zhang et al., 2007). Both Sog and Dpp bind the *Drosophila* collagen IV proteins Viking and Dcg1 (Wang et al., 2008) and modelling demonstrates that this could increase the affinity of Sog for Dpp/Scw by 50- to 100 fold (Umulis et al., 2009). Tsg releases Sog and Dpp from collagen IV (Sawala et al., 2012; Wang et al., 2008); however it is possible that, prior to release, the avidity-driven effects of Tsg are needed to achieve the ∼1000 fold increase in affinity of Sog for Dpp/Scw to permit shuttling *in vivo*.

Tsg has a well-documented role in promoting Sog/Chordin cleavage by Tld (Larraín et al., 2001; Scott et al., 2001; Shimmi and O’Connor, 2003; Xie and Fisher, 2005). However, a Sog/Chordin-independent pro-BMP activity of Tsg has been described in *Tribolium* (Nunes da Fonseca et al., 2010) and epistatic backgrounds in *Drosophila* (Wang and Ferguson, 2005), and zebrafish (Xie and Fisher, 2005). It has been proposed that this activity could be through Tsg promoting BMP ligand-receptor interactions (Berni et al., 2023; Nunes da Fonseca et al., 2010; Wang and Ferguson, 2005). Tsg binds the BMP ligand at the type I receptor (BMPRI)-binding wrist epitope (Kirsch et al., 2000a; Malinauskas et al., 2024), whereas the type II receptor (BMPRII) binds to BMP’s knuckle epitope (Kirsch et al., 2000b). Therefore, we speculate that Tsg could promote the formation of a transient Tsg-BMP-BMPRII complex. BMP receptors form heteromeric signalling complexes through an avidity mechanism (Groppe et al., 2008), therefore BMPRI would rapidly displace Tsg to form a BMPRI-BMPRII-BMP complex that activates the BMP signalling cascade (Fig. 6). As we cannot detect an interaction between Tsg and type I or II BMP receptors (Fig. S8), we propose that Tsg may aid formation of the Tsg-BMP-BMPRII complex by helping to solubilise BMP (Groppe et al., 1998), via a Tsg-BMP complex in the absence of Sog (Mi et al., 2015).

**Figure 6.**
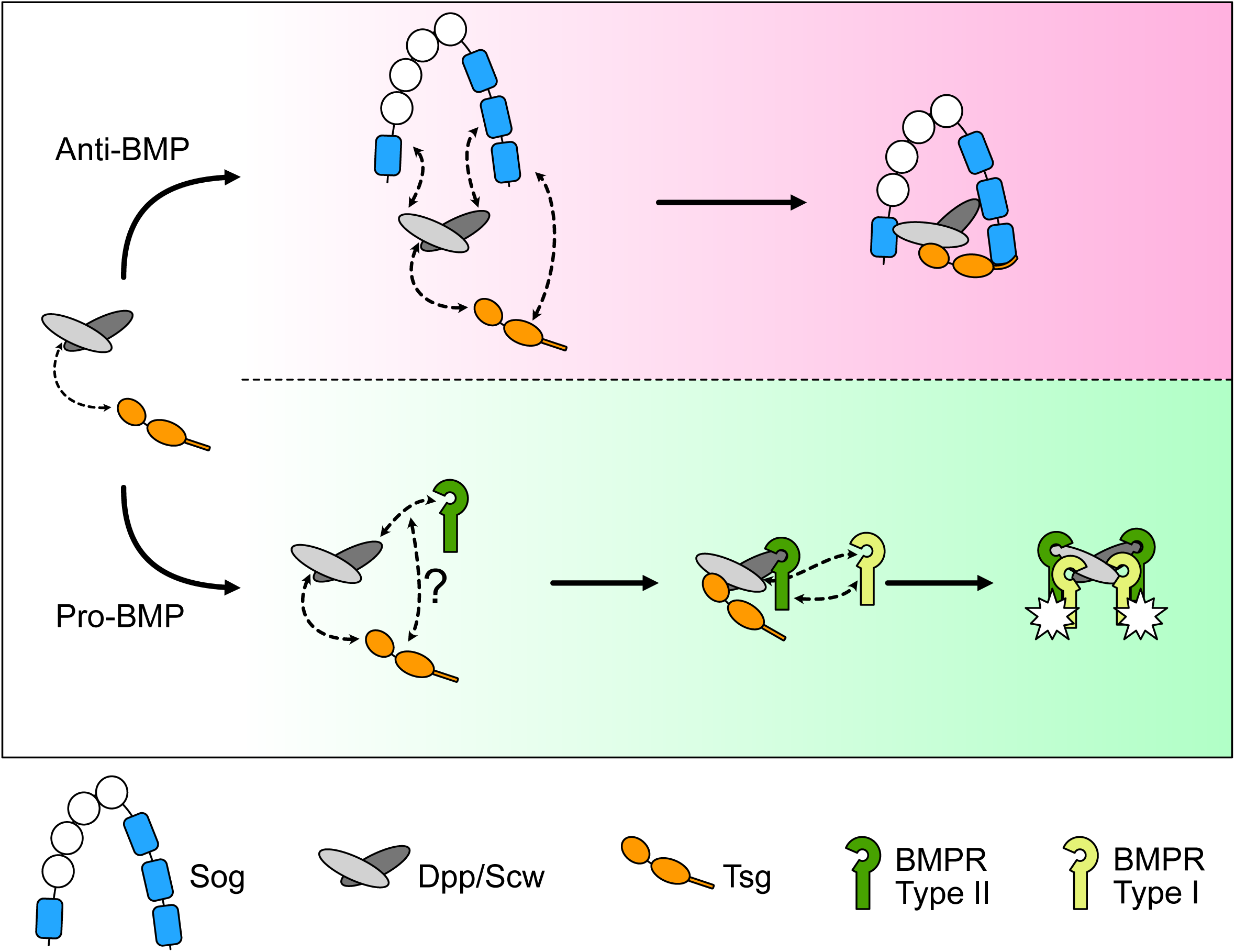
Competing avidity-based mechanism of Tsg action in extracellular BMP regulation. Top: Dpp/Scw and Sog can bind each other through two available binding sites. Tsg binds both Sog and Dpp/Scw through different binding sites to promote ternary complex formation through an avidity effect. Tsg is depicted as binding only to vWC4 of Sog throughout for simplicity but may bind other vWC domains. Bottom: Tsg promotes BMP-BMPRII interaction which rapidly assembles into a heterotetrameric BMP ligand-receptor complex, displacing Tsg, to activate the signalling cascade.

If this speculative model to explain the pro-BMP function of Tsg is correct, it suggests that Tsg can act as a catalyst to promote, through competing avidity mechanisms, either a BMP-antagonist or BMP-receptor complex (Fig. 6). We suggest that this could explain the considerable variation in Tsg function seen in insects as, by tuning the interactions of Tsg within these competing complexes, the dominating action of Tsg could be modulated. This model can explain why in *Nasonia* (Özüak et al., 2014), where *sog* is absent, Tsg displays a pro-BMP function, as Tsg would promote BMP-BMPR interaction. In *Oncopeltus*, *tsg* knockdown dorsalises the embryo consistent with an anti-BMP function (Sachs et al., 2015), but this also results in loss of *sog* expression thereby generating in effect a *tsg sog* double mutant. Some experimental evidence and modelling data suggest that there is weaker BMP signalling in this *tsg* (plus *sog)* mutant background compared to only *sog* knockdown (Sachs et al., 2015). This is suggestive of an additional Sog-independent, pro-BMP function of Tsg in *Oncopeltus*, as seen with similar epistatic backgrounds in *Drosophila* (Wang and Ferguson, 2005). In *Tribolium*, despite Sog being present, the major function of Tsg is a Sog-independent, pro-BMP function (Nunes da Fonseca et al., 2010). We note that there is poor conservation of the collagen IV binding sites in *Tribolium* Sog, so one possibility is that a weak Sog-collagen IV interaction tips the balance away from Sog-Tsg-BMP towards Tsg-BMP-BMPRII. Additional studies are required to test whether Tsg performs its opposing functions in dorsoventral patterning by, as suggested here, promiscuously participating in both agonistic and antagonist avidity-driven complexes in the extracellular space.

In conclusion, we identify the Chordin family binding site in Tsg and combine *in vitro* binding studies and *in vivo* genetics to propose an avidity-based mechanism by which Tsg promotes the formation of ternary Tsg-Sog/Chordin-BMP complexes. This furthers our understanding of this key regulatory complex across vertebrate and invertebrate development. We expect this mechanism to occur in other contexts where BMP signalling is regulated by Tsg and Chordin family members. For example in early (Graf et al., 2001; Petryk et al., 2004; Wills et al., 2006) and postnatal (Sun et al., 2010) brain development, digit patterning (Lorda-Diez et al., 2013), and colorectal cancers where both Tsg (Gylfe et al., 2013) and CHRDL2 (Sun et al., 2016) are implicated in BMP signalling misregulation.

## Materials and Methods

### AlphaFold Multimer

To perform *in silico* protein structure prediction, AlphaFold-Multimer (V2.1.1) (Evans et al., 2022; Jumper et al., 2021) was implemented on the High-Performance Computing (HPC) cluster through the Computational Shared Facility 3 (CSF3), University of Manchester. Jobs were run using 1 Graphics Processing Units (GPU) with 8 cores. The model was trained on Protein Data Bank (PDB) files deposited before 01/11/2021 and used the ‘full genetic database’ for multiple sequence alignment (MSA) configuration. Each prediction was performed with the default number of recycles to generate five relaxed predictions which were ranked based on the predicted template alignment (pTM) and interface predicted template alignment (ipTM) score defined as: 0.2 ∗ *pTM* + 0.8 ∗ *ipTM*. Predicted alignment error (PAE) plots were generated for each model through a python script (freely available: https://zenodo.org/records/15880546) implemented on the CSF3. All five model outputs were analysed, and the highest confidence prediction presented.

### Cloning, expression and purification of human CHRDL2 and wildtype and mutant human Tsg

A synthetic sequence of human Twisted gastrulation (residues Cys26-Phe223) codon optimised for mammalian expression (ThermoFisher Scientific) with a C-terminal TwinStrep-tag (IBA LifeScience) joined by a 3xGlyGlyGlySer linker was inserted into the pHLsec vector (Addgene_99845) between AgeI and KpnI restriction sites. Site directed mutagenesis to introduce single point mutations was performed by PCR and mutated PCR products annealed using In-Fusion cloning (Takara Bioscience). The human CHRDL2 sequence (Arg24-Thr429) with a C-terminal TwinStrep-tag (IBA LifeScience) joined by a 3xGlyGlyGlySer linker was also inserted into the pHLsec vector (Addgene_99845) between AgeI and KpnI restriction sites. pHLsec-Tsg-TwinStrep and pHLsec-CHRDL2-TwinStrep plasmids were transfected to Expi293-F cells (ThermoFisher Scientific) using the ExpiFectamine 293 Transfection Kit (ThermoFisher Scientific). Cells were cultured for 4 days before conditioned media was collected, centrifuged at 4500 x g at 4°C for 15 mins and filtered using 11 μm filter paper (Whatman) and then 0.65 μM PVDF filter membrane (Millipore). Recombinantly expressed protein was purified by affinity chromatography using Strep-Tactin XT 4Flow High-Capacity resin (IBA LifeSciences) and protein eluted in elution buffer (20 mM HEPES, pH 7.4, 500 mM NaCl, 75 mM Biotin). Eluted protein was further purified by size exclusion chromatography (SEC) using a Superdex 200 Increase GL column (Cytiva) on an Äkta Purifier FPLC (Cytiva) with SEC buffer (20 mM HEPES, 150 mM NaCl, pH 7.4). Protein was resolved at 0.5 ml/min, at room temperature. Protein elution from the column was monitored using UV absorbance at 280 nm and eluted in 0.5 ml fractions. Fractions of interest were assessed for protein content by SDS-PAGE and stored at −80°C. Purified, recombinantly expressed ΔN-Chordin was prepared as described previously (Troilo et al., 2014). Plasmids used are available upon request. All primers used in this study are listed in Table S2.

### Surface Plasmon Resonance

To determine binding kinetics and affinities, surface plasmon resonance (SPR) was performed on a Biacore T200 (Cytiva). Purified, recombinantly expressed ΔN-Chordin for binding studies was provided by Dr Rana Dajani (University of Manchester). Wildtype or mutant Tsg, or CHRDL2 ligand was diluted to a concentration of 30 μg/ml in sodium acetate buffer pH 5.5 and immobilised onto CM5 Series S Sensor Chips (Cytiva) via amine coupling using NHS/EDC cross linking, with exposed activated amines subsequently blocked with ethanolamine. Wildtype and mutant Tsg proteins were immobilised to 350 RU, CHRDL2 was immobilised to either 1200 RU or 3700 RU to achieve sensitive readings with both across wildtype and mutant samples. A blank reference flow cell was also activated and blocked for background subtraction. Assays were performed in running buffer (20 mM HEPES, 150 mM NaCl, 0.05% Tween20, pH 7.4) at 25°C, and all analytes were diluted in running buffer. The surface was regenerated between cycles with 0.1 M glycine pH 2 followed by 1 M NaCl pH 7.4, each at 30 μl/min for 30 secs. For kinetic assays, analytes were diluted in a two-fold concentration series and flowed over immobilised ligand at 30 μl/min for 200 secs, followed by a 300 sec dissociation phase. All data were background subtracted using the control reference surface. Binding parameters were calculated using non-linear fitting of a Langmuir 1:1 binding model performed using the Biacore T200 Evaluation Software (V2). For measurements of Tsg-BMPR1A and Tsg-BMPR2 interactions, measurements were performed on a Biacore 1K+ (Cytiva) in running buffer. A CM5 chip was immobilised with human Tsg in immobilisation buffer 0.1 M Glycine pH 4.5 to 300 RU using the standard amine coupling method. A blank flow cell was also activated and blocked for reference subtraction. Human BMPR1a (R&D Systems), BMPR2 (Abcam) -FC fusion proteins and CHRDL2 were flowed over the human TSG and blank flow cells at a concentration of 1µM for 30 seconds. Responses shown are reference subtracted.

### Biolayer Interferometry

All BLI measurements were performed on an Octet Red96 instrument (Sartorius Stedim) in an assay buffer of HBS pH 7.4 with 0.05% Tween-20. In brief, human BMPR1a (R&D systems) or human BMPR2 (Abcam)-FC chimera proteins were captured to saturating levels on hydrated Protein A sensors. A further control reference sensor was left blank. Following baseline measurements, all sensors were then dipped in a 1 µM solution of human Tsg for approximately 50 seconds and responses were recorded and reference subtracted.

### Circular dichroism

To determine secondary structure composition of wildtype and mutant human Tsg, circular dichroism (CD) was performed on a J810 spectrophotometer (JASCO). To minimise background signal from buffer, all proteins were buffer exchanged to 5 mM Tris, 150 mM NaCl pH 7.4 buffer, using a 5 ml HiTrap Desalting Sephadex G-25 column (Cytiva). Following buffer exchange, protein concentration was determined by absorbance at 280 nm on a Nanodrop-1000 spectrophotomer (ThermoFisher Scientific) and adjusted to 0.3-0.4 mg/ml. Spectra from 260-190 nm range were collected in 0.2 nm steps, with a 0.5 sec exposure time to a total of 10 accumulations which were averaged. The same measurements were taken with a blank buffer sample which was subtracted from experimental readings. Data analysis was performed with the online DichroWeb suite using the CDSSTR ((Manavalan and Johnson, 1987; Sreerama and Woody, 2000), Contin (van Stokkum et al., 1990), K2D (Andrade et al., 1993) and Selcon3 (Sreerama and Woody, 1993; Sreerama et al., 1999) algorithms with the reference set SP175t (Lees et al., 2006).

### Static Light Scattering

Protein thermal stability was assessed by static light scattering using an Uncle (Unchained Labs). Samples were diluted to 200 μg/ml in 20 mM HEPES, 150 mM NaCl, pH 7.4. Sample aggregation temperature (T_agg_) was assessed by static light scattering recorded at 473 nm through a temperature ramp from 20-95°C, in 1°C steps per min. All measurements were recorded in triplicate.

### S2R+ cell co-immunoprecipitation assay

S2R+ cells (DGRC Stock 150; https://dgrc.bio.indiana.edu//stock/150; RRID:CVCL_Z831) were cultured in Schneider’s Drosophila media (Gibco) supplemented with 10% v/v FBS, 1% v/v penicillin/streptomycin at 25°C. Cells were transfected using Effectene (Qiagen) in 12-well plates. A pMT-BiP-Tsg:ALFA plasmid was generated by PCR amplification of the Tsg coding sequence from pMT-BiP-Tsg:His (Winstanley et al., 2015) to include a C-terminal ALFA tag (Götzke et al., 2019) and inserted in pMT-BiP-V5-His (Thermofisher) digested with EcoRI and AgeI. A single Leu100Ala point mutation was introduced by In-Fusion cloning (Takara Bioscience) to generate a pMT-BiP-Tsg^L100A^:ALFA plasmid. The tail sequence (Asp209-Ser249) was excised from Tsg by infusion cloning to generate a pMT-BiP-Tsg^Δtail^:ALFA plasmid. Sog, Tsg, Tsg^L100A^ or Tsg^Δtail^ conditioned media was produced by transfecting cells with 1 μg of pMT-Sog:Myc (Sawala et al., 2012), pMT-BiP-Tsg:ALFA, pMT-BiP-Tsg^L100A^:ALFA, pMT-BiP-Tsg^Δtail^:ALFA, or pMT-BiP-Tsg^L100A,Δtail^:ALFA. Dpp/Scw conditioned media was generated as in Sawala et al., (2012) by transfecting cells in a T75 flask (Corning) with 10 μg pMT-Dpp:HA and 4 μg pMT-Scw:Flag. For the comparative Tsg^L100A,Δtail^:ALFA experiment, recombinant Dpp (Cat#159-DP, R&D Systems) was used at 100µg/mL concentration. Protein expression was induced 24 hours after transfection by supplementing media with 0.5 mM CuSO_4_. Conditioned media was collected a further 72 hours after induction and centrifuged at 5000 x g, 15 mins at room temperature. Conditioned media was then mixed and incubated at room temperature for 3 hours before binding anti-ALFA Selector PE resin (NanoTag Biotechnologies, Cat#N1510, RRID:AB_3075990) for 1 hour at 4°C. Matrix was collected and washed four times with buffer (20 mM HEPES, pH 7.4, 150 mM NaCl) and samples eluted by boiling in NuPAGE LDS Sample Buffer (ThermoFisher Scientific) + 5% β-mercaptoethanol. Samples were analysed by Western blot using Mouse anti-Myc (1:1000, Millipore, RRID: AB_11211891), Rabbit anti-ALFA (1:1000, NanoTag Biotechnologies, RRID:AB_3075998) and Mouse anti-HA (1:1000, Roche, RRID: AB_514505) primary antibodies, with IRDye® 680RD donkey anti-Rabbit IgG (1:10,000, LI-COR, RRID: AB_2716687) and IRDye 800CW donkey anti-Mouse IgG (1:10,000, LI-COR, RRID: AB_621847) secondary antibodies. Western blots were visualised on a LI-COR Odyssey CLx. Plasmids used are available upon request.

### *Drosophila* stocks and maintenance

Fly stocks were maintained on standard fly food media (yeast 50 g/l, glucose 78 g/l, maize 72 g/l, agar 8 g/l, 10% nipagen in ethanol 27 ml/l and propionic acid 3 ml/l). *D. melanogaster* stocks used in this work were: *nos-Cas9* (RRID:BDSC_78782), *tsg^attP^/FM7c-ftz-lacZ*; *tsg:ALFA* (Malinauskas et al., 2024); *tsg^L100A^:ALFA* and *tsg^Δtail^:ALFA/FM7c-ftz-lacZ* (see below). The *tsg^attP^* and *tsg^Δtail^:ALFA* stocks were balanced with *FM7c-ftz-lacZ* allowing visualisation of mutant embryos based on the absence of *lacZ* staining. *nos-Cas9* flies were used as the wildtype control throughout.

### Generation of mutant *tsg Drosophila* stocks

Both the *tsg^L100A^:ALFA* and *tsg^Δtail^:ALFA* fly stocks were generated using ΦC31-mediated transgenesis in a *tsg^attP^* stock, as outlined in Malinauskas et al., (2024). Briefly, the *tsg^attP^* stock replaces the *tsg* locus with an attP landing site. Reintegration using a RIV^white^-Tsg:ALFA plasmid repairs the *tsg* locus with all excised residues, including the wildtype *tsg* coding sequence with the addition of a C-terminal ALFA tag and a downstream *mini-white* sequence (Malinauskas et al., 2024). To generate the point mutant reintegration vector, Leu100 was mutated to Ala in the RIV^white^-Tsg:ALFA plasmid using In-Fusion cloning. To generate the truncated *tsg* sequence, the tail sequence (Asp209-Ser249) was excised from RIV^white^-Tsg:ALFA, leaving a truncated coding sequence (Met1-Glu208). Reintegration plasmids were co-injected with a ΦC31 encoding plasmid into *tsg^attP^* embryos and survivors crossed back to the balanced *tsg^attP^/FM7c-ftz-lacZ* stock. Successful recombinants were screened for using the *mini-white* marker and confirmed by sequencing genomic DNA. Male *tsg^Δtail^:ALFA* flies were not viable and so the line remained balanced over the *FM7c-ftz-lacZ* chromosome. Male *tsg^L100A^:ALFA* flies were viable and so the balancer chromosome was crossed out, to generate a homozygous *tsg^L100A^:ALFA* stock. *Drosophila* stocks are available upon request. All primers used in this study are listed in Table S2.

### smFISH imaging of *Drosophila* embryos

*Drosophila* embryos (2-4 hours) were fixed as previously described (Hoppe et al., 2020) and stored in methanol at −20°C long term. For smFISH, embryos were stained with *ush* Stellaris (Hoppe et al., 2020) and *lacZ* Stellaris (Frampton et al., 2022) probes and nuclei were stained with DAPI (1:1000, NEB 4083). Samples were mounted in ProLong Diamond antifade mountant (Thermo Fisher) and set overnight before long term storage at −20°C.

Embryos were imaged on an Andor Dragonfly200 spinning disk inverted confocal microscope with both a 20x/0.75 MImm Plan Fluor and 40x/1.3 Super Fluor objective. Samples were excited using 405 nm (5%, 100ms exposure), 561 nm (10%, 100ms exposure) and 637 nm (10%, 150ms exposure) diode lasers and collected through blue (450nm), red (600nm) or far red (700nm) filters. Each channel was collected sequentially with 4x averaging on the iXon EMCCD (1024 × 1024) camera and multiple Z slices were obtained at system optimised spacing. Images were deconvolved using inbuilt Andor deconvolution software. *tsg^Δtail^:ALFA* hemizygous males were identified by a lack of a *lacZ* signal in the 561 nm channel.

### FISH imaging of *Drosophila* embryos

Embryos (0-4h) were collected and stained by RNA *in situ* hybridisation with *ush*-digoxygenin-UTP, *lacZ*-digoxygenin-UTP, and *Race*-biotin-UTP as described (Hoppe *et al*., 2020; Kosman *et al*., 2004). Antibodies used were mouse anti-biotin (1:500, SAB4200680, Sigma-Aldrich), sheep anti-digoxygenin Fab fragments antibody, AP conjugated (1:200, Roche, cat. #11093274910 RRID: AB_514497), donkey anti-mouse IgG secondary antibody, Alexa Fluor 488 (1:500, Thermo Fisher Scientific, cat. #A-21202, RRID: AB_141607), and donkey anti-sheep IgG secondary antibody, Alexa Fluor 555 (1:500, Thermo Fisher Scientific, cat. #A-21436, RRID: AB_2535857). For pMad immunostaining, anti-Smad3 (phosphoS423+S425) [EP823Y] (1:500, Abcam, cat. #ab52903, RRID: AB_882596) primary antibody and donkey anti-rabbit IgG secondary antibody, Alexa Fluor 647 (1:500, Thermo Fisher Scientific, cat. #A-31573, RRID: AB_2536183) were used. Embryos were incubated with DAPI (1:250, NEB 4083) to stain the nuclei, followed by mounting in ProLong Diamond Antifade Mountant (Thermo Fisher Scientific, P36961).

Embryos were imaged on a Zeiss 980 Axio Observer 7 with a LD LCI Plan-Apochromat 40x/1.2 Imm Korr DIC M27 FCS objective with 0.6x zoom. Confocal settings were as follows: pinhole (0.77AU/0.88AU/1AU), LSM scan speed 7, bidirectional, 4x line averaging, image size 4293×4293 pixels, and Z step size of 0.25µm. Images were collected using GaAsP-PMT detectors using the 405nm (0.2%), 488nm (1%), 561 (1%), and 639 (0.2%) laser lines. Images were processed with LSM Plus Processing (Zeiss). Images shown are maximum intensity projections, and only *tsg* embryos negative for *lacZ* present on the balancer chromosome (i.e. *tsg* mutant) were imaged.

### Image analysis

The widths of the pMad, *Race* and *ush* expression/activation domains were quantified in deconvolved images in Fiji (Schindelin et al., 2012). Maximum intensity projections of imaging stacks were made, and nuclei counted at 50% embryo length across each domain in a line perpendicular to the anterior-posterior axis. Statistical analysis was performed in GraphPad Prism 10 (v.10.2.2).

### Amnioserosa counts

*Drosophila* embryos (5-7 hours) were fixed as previously described (Hoppe et al., 2020) and stored in methanol at −20°C long term. Fixed embryos were stained with mouse anti-Hindsight 1G9 (1:40, DSHB Cat No. 1g9, RRID: AB_528278) and anti-Mouse IgG AP conjugate S372B (1:500, Promega). For *tsg^Δtail^:ALFA* embryos, homozygous mutants were identified by the absence of lacZ staining by co-staining using a chicken anti-Β-Gal (1:5000, Abcam, Cat#ab9361, RRID:AB_307210) primary antibody and anti-chicken IgY AP conjugate (1:2000, Sigma, Cat#A9171). Stage 11 embryos were imaged on a Leica DM6000 B microscope with a 40x objective using brightfield illumination. Amnioserosa cells were counted on ImageJ using the Cell Counter plugin. 50 embryos across two replicates were analysed and counts plotted, and statistical analysis performed in GraphPad Prism 10 (v.10.2.2).

### Viability assays

Embryos (0-4h) were plated onto sugar plates (1.25% agar, 3.75% w/v sucrose, 0.25% propionic acid) and incubated at 25°C. Larvae were counted at 24 hours and again at 48 hours, to ensure that all viable embryos that developed into larvae were included.

### Phylogenetic analysis of *tsg* sequences

To determine the evolutionary conservation of the *Drosophila tsg* C-terminal extension, a reference proteome was used. This reference proteome is curated by UniProt to landmark proteome space and has been selected manually and algorithmically to provide a representative overview of taxonomic diversity. The text search function was used on UniProt, queried with ‘Twisted Gastrulation and Reference Proteome’. This returned 74 entries which were manually curated to remove closely related species and poorly annotated sequences, yielding a list of 60 species sequences. This condensed list was submitted to the NCBI Taxonomy Tree browser (Schoch et al., 2020) and the results exported as a .phylip file which was then visualised and annotated in Mega11 (Stecher et al., 2020; Tamura et al., 2021).

## Supporting information

Moore et al SI

## Acknowledgments

We thank Sanjai Patel and the Manchester Fly Facility for assistance in generating the *tsg* mutant fly lines, Heather Johnson for assistance with the western blots, Dr Thomas Jowitt (University of Manchester Biomolecular Analysis Facility) for advice on the circular dichroism and SLS, Dr Rana Dajani for providing ΔN-chordin protein, and the University of Manchester Bioimaging facility.

## Author contributions

Conceptualisation, G.M., C.B. and H.L.A.; investigation, G.M., R.R., L.F.B., C.S., H.B.; writing, G.M., C.B., and H.L.A.; supervision, C.B. and H.L.A.; funding acquisition, C.B. and H.L.A.

