## Supplementary material for "An avidity-driven mechanism of extracellular BMP regulation by Twisted gastrulation": Moore et al SI

### Supplementary information

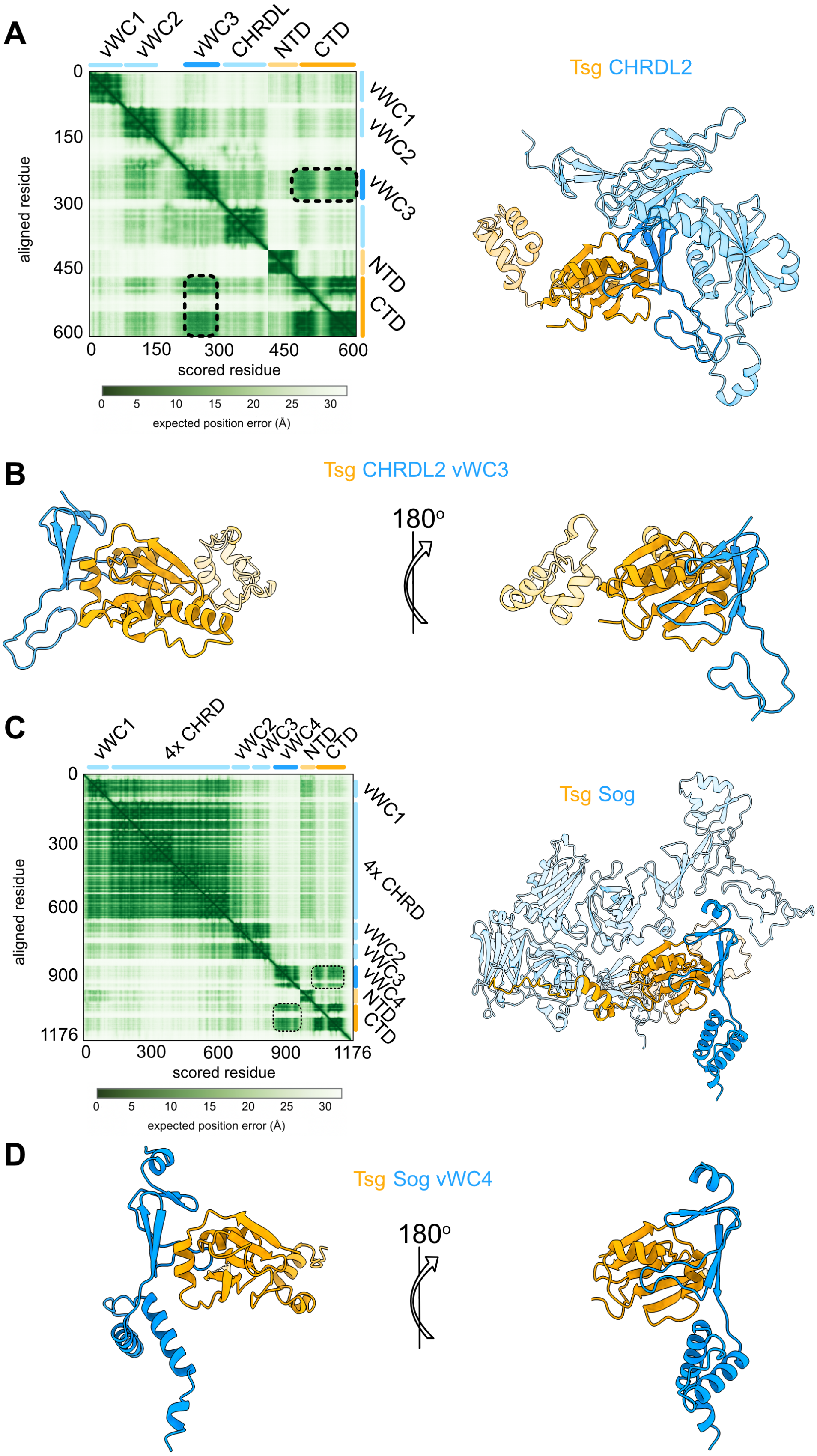


**Figure S1. AlphaFold models of Tsg and Chordin family antagonists.**
(A) AlphaFold model (right) and corresponding PAE plot (left) of human Tsg (orange) – CHRDL2 (blue) complex with a putative interface between Tsg CTD (dark orange) and CHRDL2 vWC3 (dark blue) as indicated by the dashed regions on the PAE plot. (B) AlphaFold model of Tsg-CHRDL2 vWC3 complex, surface hydrophobicity plotted. (C) and (D) as above but for *Drosophila* Tsg-Sog complex.

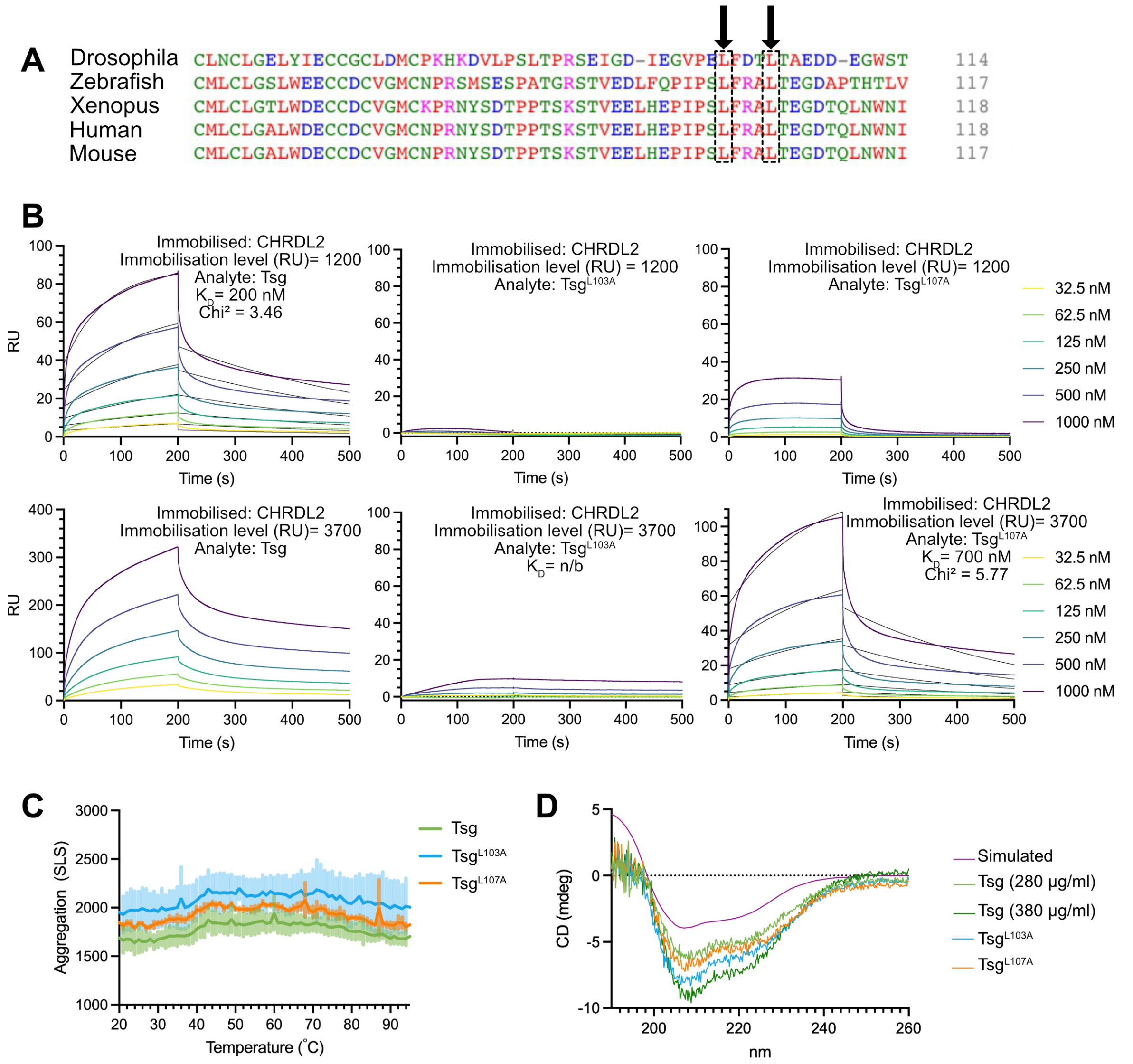


**Figure S2. Binding, stability and foldedness of Tsg point mutants.**

(A) Sequence alignment of Tsg sequences from five species showing conservation of identified Leu residues (black arrows). (B) SPR sensorgrams of human Tsg, $Tsg^{L103A}$, or $Tsg^{L107A}$ binding to immobilised CHRDL2. CHRDL2 was immobilised at two surface densities for optimal signal detection of both wild type and mutant Tsg binding. As such, binding kinetics for wild type Tsg were calculated from the low surface density measurements, and mutant Tsg kinetics calculated from the high surface density measurements. Binding kinetics were calculated by fitting curves to a Langmuir 1:1 binding model. All experiments were performed in triplicate, average curves plotted with model fit in black. n/b = no binding. (C) Static light scattering (SLS) traces of wild type Tsg (green), $Tsg^{L103A}$ (blue), and $Tsg^{L107A}$ (orange) recorded at 473 nm from 25-95°C. Mean as solid line of triplicate samples with standard deviation from the mean. (D) Buffer subtracted circular dichroism spectra of wild type Tsg at 280 µg/ml (light green) and 380 µg/ml (dark green), $Tsg^{L103A}$ at 330 µg/ml (blue) and $Tsg^{L107A}$ at 290 µg/ml (orange) with a simulated spectrum based on the crystal structure 8BWN (purple). The theoretical spectrum was generated using PDB2CD webserver from the crystal structure 8BWN of Tsg. For all samples, 10 spectra were recorded and averaged.

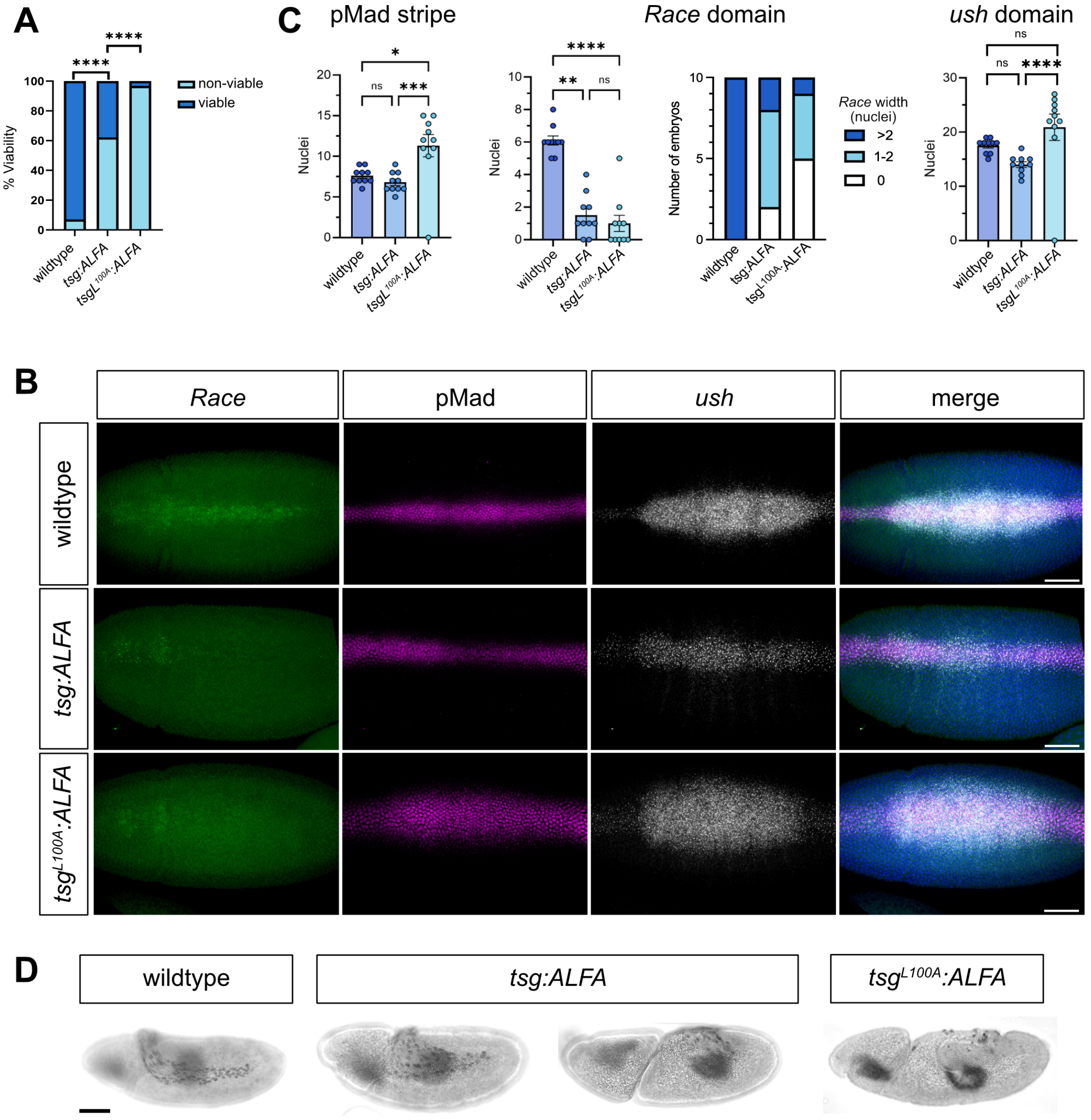


**Figure S3. Viability and patterning of *wildtype* and *tsg* embryos.**
(A) Graph shows the percentage of control wildtype, *tsg:ALFA*, or $tsg^{L100A}$*:ALFA* embryos that reach the larval stage. For *wildtype* n= 442, for *tsg:ALFA* n= 492, and for $tsg^{L100A}$*:ALFA* n= 462, across two biological repeats. Fisher's exact test: ****, $P \leq 0.0001$. (B) Fluorescence in situ hybridisation of *Race* and *ush* expression and immunostaining of pMad distribution in control, *tsg:ALFA*, or $tsg^{L100A}$*:ALFA* stage 5 embryos (dorsal views). Note that, using FISH, *Race* expression is less evident in the *tsg:ALFA* embryos compared to wildtype, whereas we previously detected a more obvious *Race* stripe in *tsg:ALFA* embryos using smFISH (Malinauskas et al., 2024). We speculate that this difference relates to the higher sensitivity of smFISH. Nuclei are stained with DAPI. Scale bar = 50 μm. (C) Quantitation of the number of nuclei expressing *Race* or *ush*, or that are pMad-positive at 50% embryo length. Kruskal-Wallis with Dunn's multiple comparison test: ns, $P > 0.05$; *, $P \leq 0.05$; **, $P \leq 0.01$; ***, $P \leq 0.001$; ****, $P \leq 0.0001$. n=10. Error bars are SEM. (D) Brightfield images of stage 11 wildtype, *tsg:ALFA*, and $tsg^{L100A}$*:ALFA* embryos stained for Hnt to mark amnioserosa cells (lateral views). Two *tsg:ALFA* embryos are included to reflect the variability in amnioserosa cell number as shown by the quantitation in Fig. 3E. Scale bar = 50 μm.

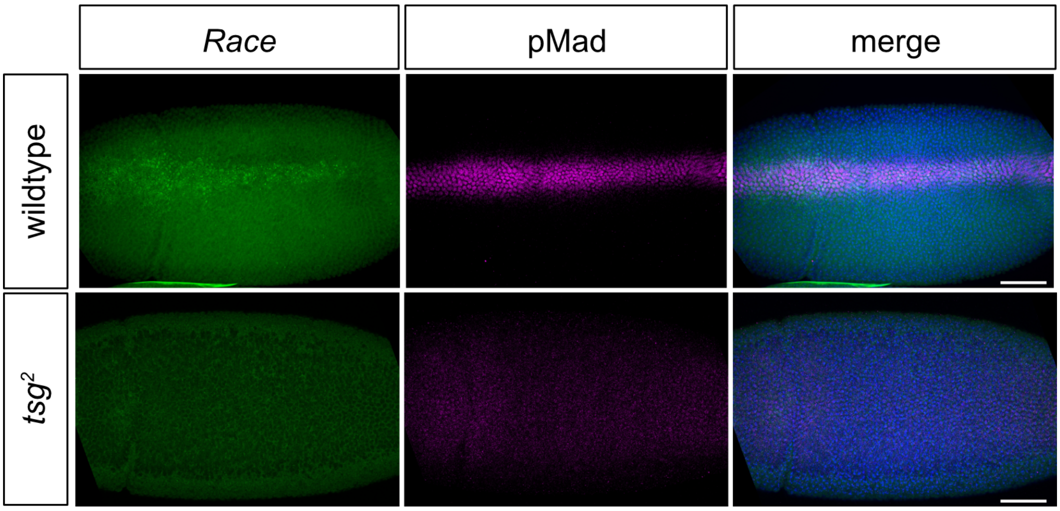


**Figure S4. *Race* expression domain and pMad pattern in *tsg*$^2$ *embryos.***
Fluorescence in situ hybridisation of *Race* expression and immunostaining of pMad in control and *tsg*$^2$ embryos (dorsal views). Nuclei are stained with DAPI. Scale bar = 50 µm.

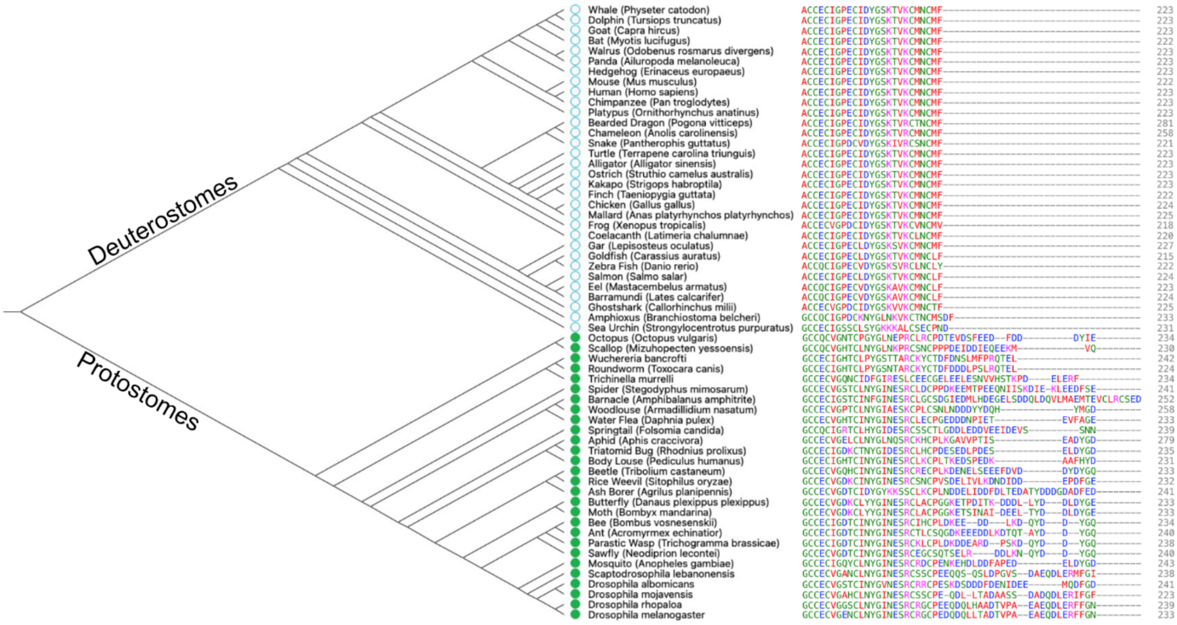


**Figure S5. Protostomes but not deuterostomes have a conserved C-terminal Tsg tail sequence.**
Straight cladogram and accompanying sequence alignment showing the evolutionary divergence of the Tsg tail, with deuterostomes lacking (blue circle) and protostomes conserving (green dot) a Tsg tail. Cladogram rooted and not to scale.

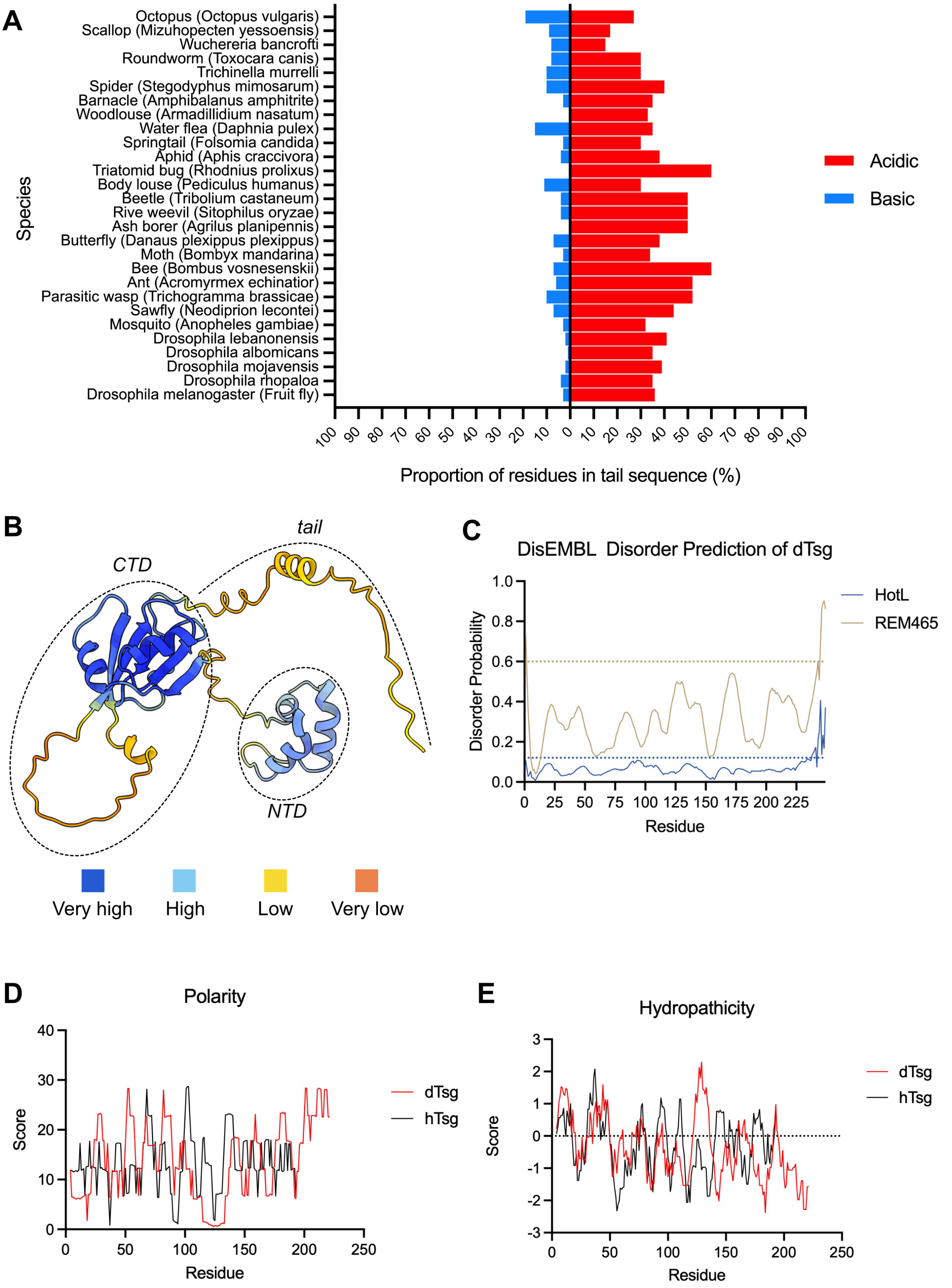


**Figure S6.** ***In silico*** **tools predict a disordered Tsg tail.**
(A) Percentage of residues in the Tsg tail sequences that are acidic (Asp and Glu) or basic (Arg and Lys) by species. (B) AlphaFold model of *Drosophila* Tsg predicts the N-terminal domain (NTD) and C-terminal domain (CTD) with very high (pLDDT >90) or high

(90>pLDDT>70) confidence but the tail sequence with low (70>pLDDT>50) or very low (pLDDT<50) confidence. pLDDT, predicted Local Distance Difference Test, plotted per colour key as shown below. AlphaFold identifier: AF-P54356-F1. (C) DisEMBL disorder prediction algorithms HotLoops and REMARK-465 for the *Drosophila tsg* sequence, dotted line represents model random expectation threshold. The Tsg tail (D) increases the overall polarity score (Zimmerman et al., 1968) and (E) decreases the overall hydropathicity score (Kyte and Doolittle, 1982) of *Drosophila* Tsg (dTsg), with human Tsg (hTsg) for comparison.

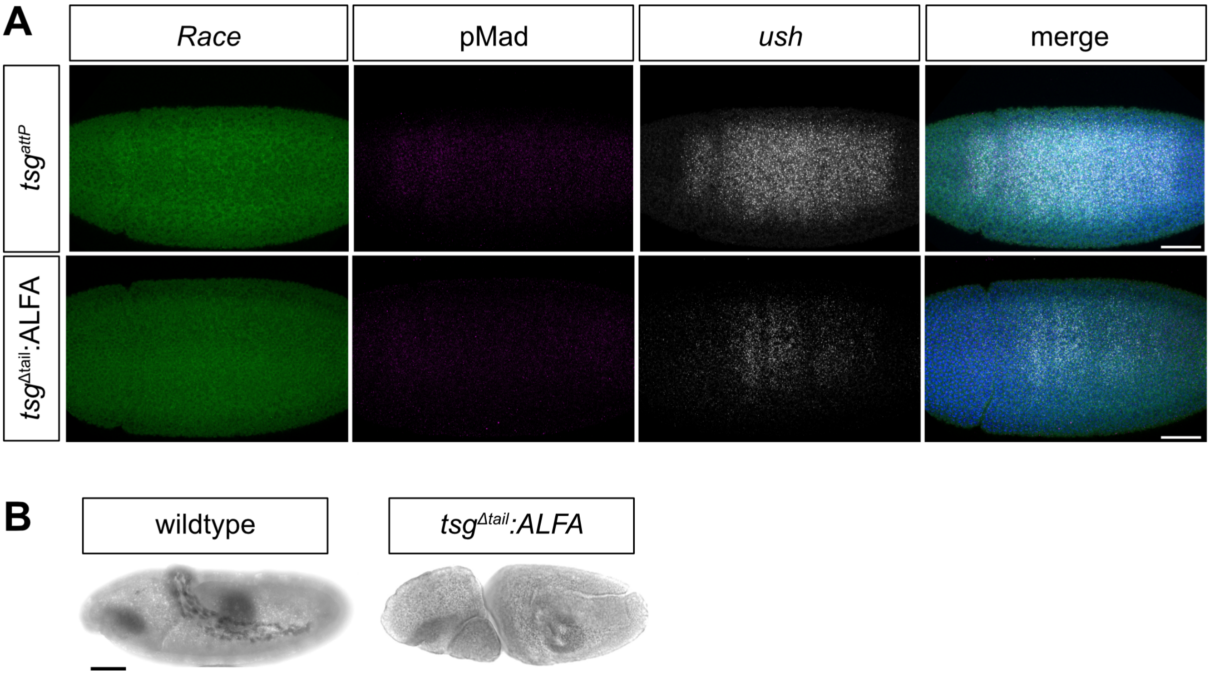


**Figure S7. Patterning and amnioserosa staining of *tsg* mutant embryos.**
(A) Fluorescence in situ hybridisation of *Race* and *ush* expression domains and immunostaining of pMad in $tsg^{attP}$ and $tsg^{\Delta tail}$*:ALFA* embryos (dorsal views). Nuclei are stained with DAPI. Scale bar = 50 μm. (B) Brightfield images of *wildtype* and $tsg^{\Delta tail}$*:ALFA* at stage 11, stained for Hnt to mark amnioserosa cells (lateral view). Scale bar = 50 μm.

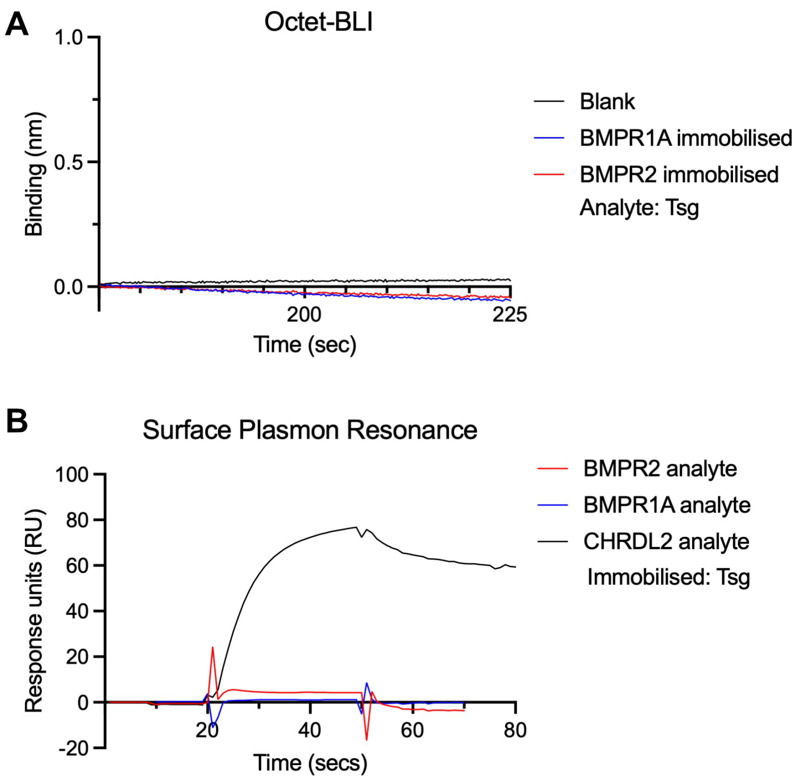


**Figure S8. Tsg does not bind BMPR1A or BMPR2 in biophysical assays.**
(A) Octet-BLI measurements of binding between BMPR1A or BMPR2 (immobilised) with human Tsg (analyte). (B) SPR sensorgrams of either CHRDL2 (control), BMPR1A or BMP2 flowed over immobilised human Tsg. NB: spikes in traces due to injection of analyte.

**Table S1. Estimated secondary structural element proportions in Tsg wildtype and mutants from circular dichroism measurements.**

Tsg (280 μg/ml)

| Method | α-Helix | β-strand | Turns | Unordered | NMRSD |
|---|---|---|---|---|---|
| **Contin** | 0.055 | 0.182 | 0.159 | 0.467 | 0.165 |
| **CDSSTR** | 0.07 | 0.24 | 0.14 | 0.41 | 0.094 |
| **K2D** | 0.14 | 0.33 | n/a | 0.53 | 0.627 |
| **Selcon3** | 0.051 | 0.169 | 0.13 | 0.434 | 0.278 |
| **Mean** | **0.079** | **0.23025** | **0.143** | **0.46025** | **n/a** |

Tsg (380 μg/ml)

| Method | α-Helix | β-strand | Turns | Unordered | NMRSD |
|---|---|---|---|---|---|
| **Contin** | 0.052 | 0.19 | 0.157 | 0.46 | 0.099 |
| **CDSSTR** | 0.07 | 0.23 | 0.13 | 0.42 | 0.059 |
| **K2D** | 0.14 | 0.33 | n/a | 0.54 | 0.577 |
| **Selcon3** | 0.053 | 0.113 | 0.137 | 0.395 | 0.132 |
| **Mean** | **0.07875** | **0.21575** | **0.14133333** | **0.45375** | **n/a** |

$Tsg^{L103A}$

| Method | α-Helix | β-strand | Turns | Unordered | NMRSD |
|---|---|---|---|---|---|
| **Contin** | 0.066 | 0.169 | 0.148 | 0.499 | 0.165 |
| **CDSSTR** | 0.07 | 0.23 | 0.14 | 0.42 | 0.061 |
| **K2D** | 0.13 | 0.33 | n/a | 0.54 | 0.579 |
| **Selcon3** | 0.066 | 0.155 | 0.14 | 0.426 | 0.262 |
| **Mean** | **0.083** | **0.221** | **0.14266667** | **0.47125** | **n/a** |

$Tsg^{L107A}$

| Method | α-Helix | β-strand | Turns | Unordered | NMRSD |
|---|---|---|---|---|---|
| **Contin** | 0.062 | 0.169 | 0.161 | 0.47 | 0.144 |
| **CDSSTR** | 0.07 | 0.23 | 0.14 | 0.41 | 0.07 |
| **K2D** | 0.14 | 0.32 | n/a | 0.55 | 0.544 |
| **Selcon3** | 0.056 | 0.17 | 0.136 | 0.441 | 0.224 |
| **Mean** | **0.082** | **0.22225** | **0.14566667** | **0.46775** | **n/a** |

#### Table S2 Primers used in this study

| Primer name | Sequence |
|---|---|
| dTsg L100A Fwd | GCCCGAGGCCTTCGACACCCTAACCGCCG |
| dTsg L100A Rev | TCGAAGGCCTCGGGCACTCCTTCAATGTC |
| dTsg_Δtail_Fwd | GCCGCGAGATCTCGTCGAGAATGCAGTTACTGTGCTACTTCGTCA |
| dTsg_Δtail_Rev | CGCCGCCCTCCGGACAACCGCGGCAGC |
| dTsg Sequencing Fwd | GCCATCTACCTGACCATCTGGATAC |
| dTsg Sequencing Rev | TGGATCCACTGTTAGAACTAAAAGG |
| hTsgL107A_Fwd | CAGAGCCGCCACCGAGGGCGATACCCAGC |
| hTsgL107A_Rev | TCGGTGGCGGCTCTGAACAGGCTAGGAATGG |
| hTsgL103A_Fwd | TCCTAGCGCCTTCAGAGCCCTGACCGAGG |
| hTsgL103A_Rev | CTGAAGGCGCTAGGAATGGGCTCGTGC |
